# DNA Polymerase α has pyrimidine dimer translesion activity that is suppressed during normal replication

**DOI:** 10.1101/2025.07.02.662766

**Authors:** Projit Mukherjee, Abheerup Sarker, Grant D. Schauer

**Author notes:** **Author Contribution:** P.M., A.S., and G.D.S. performed experiments and analyzed data. P.M. and G.D.S. wrote the manuscript.

## Abstract

To ensure rapid and accurate DNA replication during S-phase, the cell uses DNA damage tolerance (DDT) pathways, leaving lesions to be repaired after completion of replication. One established DDT pathway is Translesion Synthesis (TLS), in which lesions are bypassed by specialized TLS Polymerases that work in conjunction with the replisome. Here we demonstrate that DNA Polymerase alpha (Pol α), the replicative primase/polymerase, can also unexpectedly replicate through bulky lesions *in vitro*. We use biochemical and single-molecule fluorescence assays to characterize cyclobutane pyrimidine dimer (CPD) TLS activity of Pol α. We observe that Pol ε, the leading strand replicative polymerase, and RPA, a single-stranded DNA binding protein complex, both strongly inhibit CPD TLS activity of Pol α. In contrast, Pol η, the canonical TLS Pol for pyrimidine dimers, is unaffected by Pol ε and is conversely stimulated by RPA. Finally, we demonstrate with single-molecule Fluorescence Resonance Energy Transfer (FRET) that the DNA binding cleft of Pol α must remain in the open state to accommodate a bulky CPD lesion during TLS, possibly accounting for the relatively slow kinetics of CPD bypass that we observe. The results suggest that the intrinsic bulky TLS activity of Pol α is likely suppressed at the replication fork by the replisome itself during normal replication.

**RESEARCH HIGHLIGHTS:**

- Polymerase α has translesion synthesis (TLS) activity past a CPD lesion, the first demonstration of TLS activity at a bulky lesion by a replicative polymerase at physiological nucleotide levels.
- Intrinsic Pol α TLS activity is suppressed during normal replication by replisome components Pol ε and RPA.
- Pol ε suppresses TLS incorporation by Pol α by proofreading
- The binding cleft of Pol α stays open to accommodate a pyrimidine dimer.

**GRAPHICAL ABSTRACT:** Pol α harbors CPD TLS activity that is suppressed by RPA and Pol ε at the replication fork during normal replication.

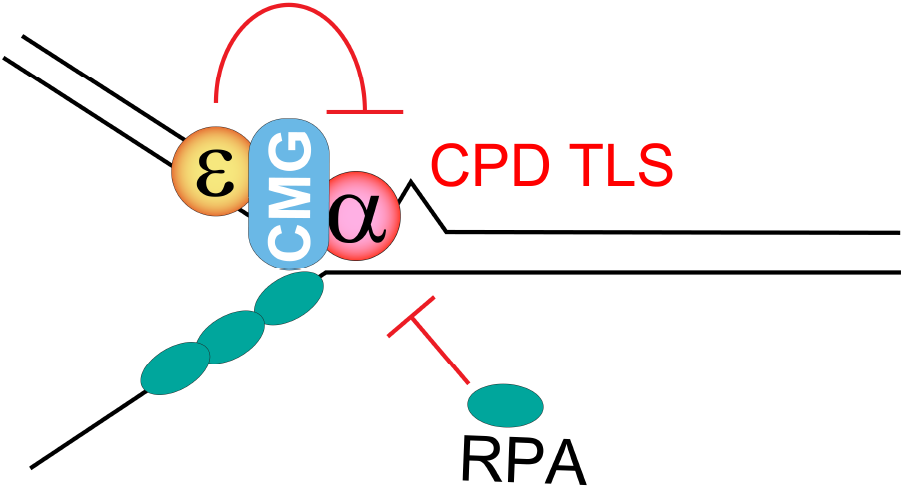

## INTRODUCTION

Genomes are copied quickly during S-phase to avoid potentially mutagenetic or fatal replication cataclysms [1]. In budding yeast, the core replication machinery (a.k.a. replisome) consists of the 11-subunit CMG helicase (cdc45, Mcm2-7, GINS), Polymerase (Pol) α, a hetrotetramer with both primase and polymerase activity that initiates hybrid RNA/DNA primers *de novo* on parental DNA, Pol ε (a heterotetramer), Pol δ (a heterotrimer), RFC (Replication factor C) a heteropentameric clamp loader, the homotrimeric PCNA (Proliferating Cell Nuclear Antigen) sliding clamp, and the heterotrimeric RPA (Replication Protein A), which coats exposed single-stranded DNA [2]. The leading and lagging strand Pols (Pol ε and Pol δ, respectively) are very accurate due 5’-3’ exonuclease (i.e., proofreading) activity, which is not present in Pol α. Stringency is also afforded to these Pols due to their sterically constrained DNA binding grooves; as such, active replication forks stall at bulky lesions in their path, leaving them susceptible to fork collapse and genetic instability [1–3].

Translesion synthesis is a well-characterized DNA Damage Tolerance (DDT) pathway in eukaryotic cells, whereby lesion-stalled replication forks are rescued by dedicated translesion synthesis polymerases (TLS Pols) that help the replisome bypass damage on-the-move, accurately incorporating bases opposite from the damage [1,4–9]. *S. cerevisiae* expresses three TLS Pols: Pol η, Pol ζ, and Rev1. Of these, Pol η, a Y-family monomeric polymerase, shows specificity and high accuracy in incorporating across cyclobutene pyrimidine dimers (CPDs; *a*.*k*.*a. cis-syn* thymine dimers), which are bulky dibasic lesions formed upon exposure to UV light [8,10–12]. This activity is considered to be made possible by the large and open DNA binding cleft of Pol η in both yeast and humans, which coordinates the incorporation of two deoxyadenosine (dA) bases in frame across from a thymine dimer [13,14]. Relatively error prone compared to replicative polymerases due to their permissive active sites and lack of proofreading activity, these TLS Pols are typically limited to this brief but critical task [9]. For instance, Pol η requires PCNA clamp for activity, and it has been shown that PCNA ubiquitination serves as a beacon for Pol η recruitment in both yeast and human cells, indicating that TLS activity is highly regulated to prevent excessive error-prone replication [3,6,8,15–17].

Polymerases across domains share many structural and functional similarities [18,19]. Some replicative polymerases have been demonstrated to have intrinsic TLS activity, yet, to the best of our knowledge, none have been shown to polymerize across bulky lesions like CPD except two reports demonstrating TLS activity of Pol δ at high (≥0.1 mM) levels of deoxynucleotide triphosphates (dNTPs) [20,21]. Recently, it was discovered that Pol α can easily bypass a lagging strand CPD lesion by repriming ahead of a CPD lesion (a.k.a. lesion skipping [5]), but that leading strand CPD bypass via repriming was very inefficient [10]. Separately, Primpol, a protein from the archaeal-eukaryotic primase (AEP) family that is expressed in human cells likely as a horizontal gene transfer from viruses, has been shown to play a role in regular DNA replication by not only repriming past lesions, but likely also by performing TLS activity past CPDs and various other bulky lesions with its native polymerase activity [20–24]. Inspired by these findings, we asked whether Pol α might also harbor TLS activity across bulky lesions: even though they are only distally related evolutionarily through their shared homology in their primase domains, Pol α is enzymatically related to Primpol in that they share both primase and polymerase activities and are relatively error prone due to lack of exonuclease activity. Indeed, even more recently, it was found that Pol α can bypass non-bulky lesions like thymine glycol, suggesting that a lack of proofreading may be enough to drive error-prone TLS [27]. Whether this apparent TLS activity extends to bulkier lesions that are more sterically unfavorable for bypass has not been explored.

Pol α comprises 4 subunits: Pol1 (167 kDa), the large subunit that contains the polymerase active site, and Pri1 (48 kDa), which harbors primase activity; each respectively works with regulatory subunits Pol12 (79 kDa) and Pri2 (62 kDa) to dynamically orchestrate *de novo* synthesis of RNA/DNA hybrid Okazaki fragments. Pol α is tethered to CMG helicase near the nascently unwound lagging strand, and through a series of large conformational changes, synthesizes ∼10 nt RNA primers and subsequently extends them with ∼15-20 nt DNA to make ∼25nt hybrid RNA/DNA primers [28–30]. Pol α also lacks proofreading capability and does not work with PCNA sliding clamp, and thus is the least accurate of the replicative polymerases [3]. In this report, we discover and characterize CPD TLS activity of Pol α. We observe that TLS activity can occur either on simple primed templates (i.e., lagging strand models) or working with on the leading strand with CMG helicase, incorporating dNTPs across from a pyrimidine dimer lesion. We further discover inhibitory mechanisms of this activity from replisome components like RPA and Pol ε that are potentially responsible for suppressing Pol α activity *in vivo*.

## RESULTS

### CPD bypass by Pol α on a lagging strand model template

We first performed primer extension assays on simple template/primer substrates (i.e. lagging strand models) in the absence (Ld/P) or presence (Ld^CPD^/P) of a CPD downstream of the primer (**Figure 1A**). When we replicated this template with Pol α, CPD bypass could be observed by noting the existence of a full-length product past the lesion (**Figure 1B**), demonstrating bypass at a relatively slow rate (∼2.6% bypass/min) (**Figure 1C**). Interestingly, the CPD bypass we observe is markedly slower than the recently reported Pol α TLS activity on nonbulky abasic, thymine glycol, and 8-oxo-G lesions, which reach near complete bypass efficiency after around 1 min [27]. Since we monitored extension of a radiolabeled primer in the presence of dNTPs but lacking any rNTPs, this result unambiguously demonstrates that Pol α has TLS activity via its polymerase domain *in vitro*, as this is the only way to produce full length product.

**Figure 1.**
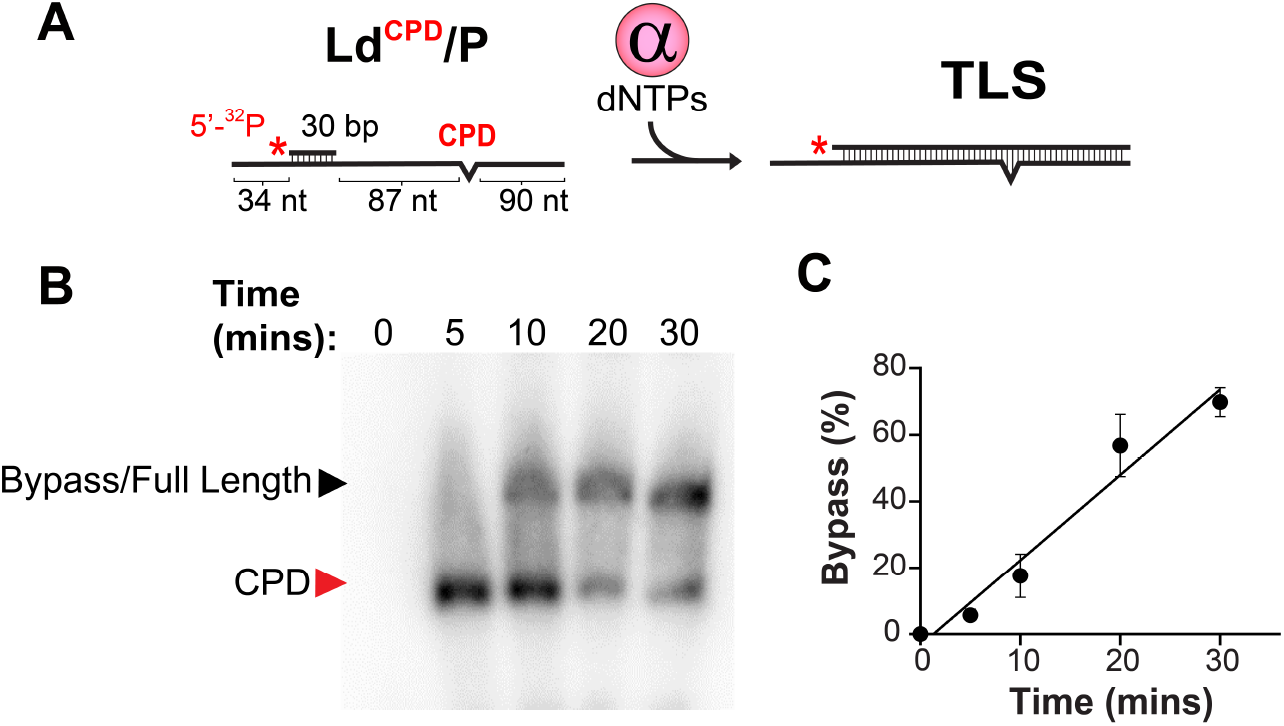
CPD bypass by Pol a on a lagging strand model. **A)** Schematic showing Pol α TLS activity on a primed template bearing a CPD lesion (Ld^CPD^/P). **B)** Representative radiolabeled primer extension assay showing CPD bypass kinetics of Pol α. Pol α exhibits TLS activity (i.e. completion to full length) in the presence of dNTPs. The location of the CPD lesion (117 nt) and the full length product (209 nt) are indicated. Reactions were stopped at the indicated times and visualized by denaturing PAGE. **C)** Quantification of bypass kinetics (2.6% bypass/min; R^2^=0.94). Data represents mean +/-STD of bypass efficiency, the ratio of full length to stalled product intensity, taken from three separate experiments.

Since it has been shown that replicative polymerases can spuriously incorporate ribonucleotides [31,32] and since misincorporation of ATP across from the fused thymine dimer could be a factor in the experiments below in which high ATP levels are required for DNA unwinding by CMG helicase, we next asked whether Pol α could demonstrate ribonucleotide misincorporation across from the CPD. Adding a limiting set of nucleotides (dGTP, dCTP, dTTP) that lacked dATP and titrating in ATP, we found that Pol α can indeed incorporate rA bases from ATP during regular polymerization, especially at 5 mM ATP, yet misincorporation was moderate and we did not observe marked synthesis up to the lesion by 30 min (**Figure S1A**). We note that we attempted to test another primer that was only 4 nt upstream of the CPD such that the only dATP incorporation possible would be across from the thymine dimer, but we could not observe primer extension in this case (not shown), indicating that Pol α may not prefer to initialize replication close to a lesion, possibly due to steric hinderance during binding. We next asked whether the ATP used for CMG helicase activity in experiments below could be responsible for CPD TLS activity. First testing the effect of 1 mM ATP or ATP analogues on Pol α TLS activity, we noted that these NTPs appeared potentially inhibitory to TLS (**Figure S1B**). Exploring this further by titrating ATP to 5 mM, we showed a dose dependent inhibition of TLS activity (**Figure S1C**, quantified in **Figure S1D**), indicating that the predominant mechanism of CPD bypass by Pol α we observe *in vitro* occurs via canonical dATP incorporation.

### CPD bypass by Pol α on the leading strand

We next asked whether Pol α CPD TLS could work in the context of an actively elongating replication fork. In these experiments, replication was reconstituted on a forked dsDNA template with pure proteins either with or without a CPD 33 bases downstream of the fork junction. In the forked substrate we use (Ld/Lg/P and Ld^CPD^/Lg/P)The leading strand templates are identical to that used in **Figure 1** and **Figure S1** but have been annealed with a 185 nt lagging strand, providing a noncomplementary fork that is required for CMG loading *in vitro*. The reaction is staged in three incubation steps [33]: CMG helicase is first loaded onto the fork, then polymerases and clamp are introduced, and the reaction is finally initiated with ATP and dNTPs (see schematic in **Figure 2A**). Without polymerase, the primer is not extended (**Figure 2A**, Lane 1). Without CMG helicase, the primer can only reach the fork junction since Pol ε or Pol α do not have strand displacement capability (**Figure 2**, Lane 2); however, Pol α appears to become primarily stalled on the template at a dT-rich region we use to promote CMG loading up the fork duplex (**Figure 2**, lane 3). When CMG is loaded, followed by Pol ε, a tight complex known as CMGE is formed, constituting a minimal leading strand replisome [34,35]. In the absence of a CPD, the replisome was able to extend the primer to full length (**Figure 2A**, Lanes 5-6).

**Figure 2.**
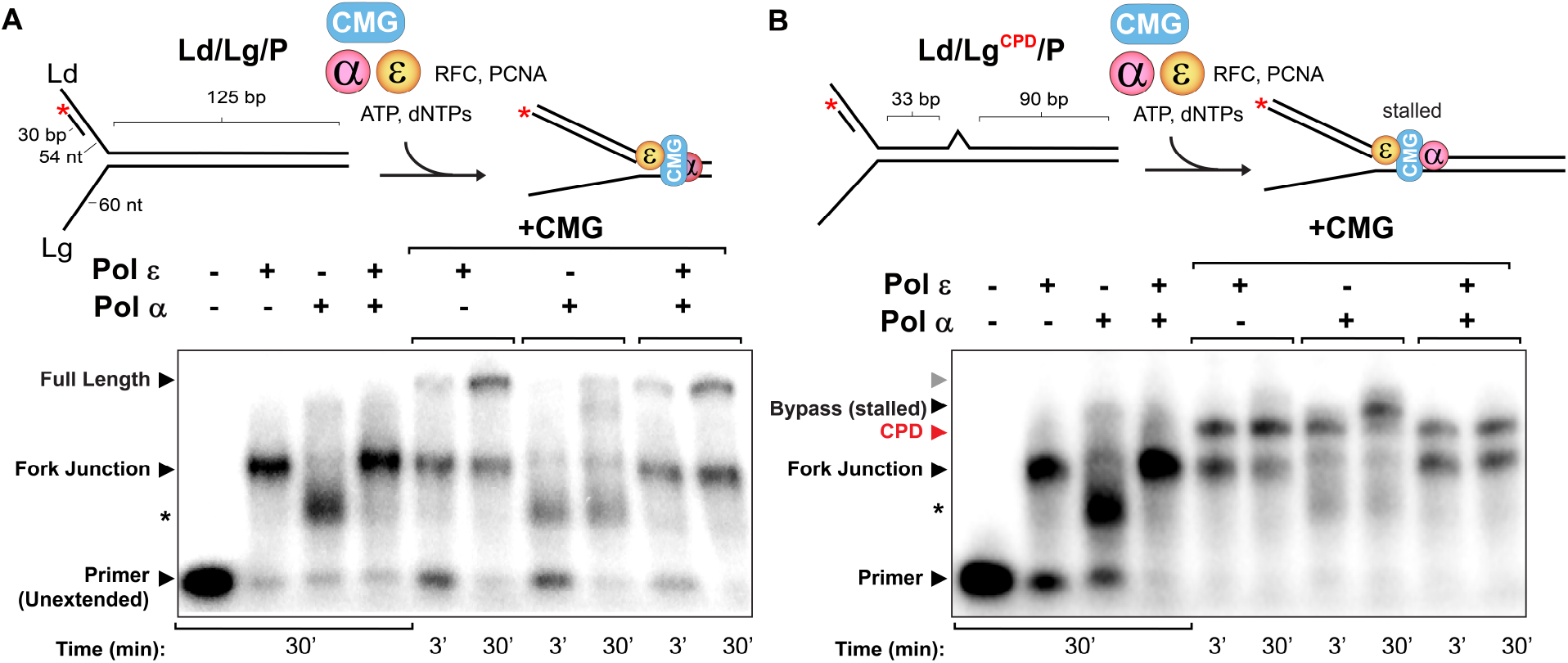
CPD bypass by Pol α on the leading strand. Primer extension assays of **A)** Replication on forked substrate without lesion and **B)** Replication on forked substrate with CPD lesion. Top (both panels): Reaction scheme. CMG is loaded first, followed by incubation with Pols α and/or ε, RFC clamp loader, and PCNA clamp, and the reaction is initiated with ATP and dNTPs. Bottom (both panels): Denaturing gel of primer extension assay. Reactions with the indicated Pols were performed. Templates were preloaded with CMG where indicated. Reactions were stopped at the times indicated. Expected product lengths are 30 nt for unextended primer, 84 nt for replication to the fork junction, 117 nt for CPD-stalled, and 209 nt for full length product. Asterisk indicates poly(dT) region where Pol α stalling is expected to occur. The gray arrow in panel B shows the length of the (missing) full length product.

However, in the presence of a mid-duplex CPD, the replisome stalls at the site of the lesion, even after 30 min (**Figure 2B**, Lanes 5-6). Here, Pol ε cannot bypass a CPD lesion, likely due to its sterically stringent active site that prevents reading through bulky lesions, which has been reported elsewhere [10,29,30]. We have previously observed that Pol α can work with CMG on the leading strand [38]. In this CMG-Pol α condition, replication was able to proceed through the CPD in a time-dependent manner—there was little observable bypass at 3 min, but by 30 min the bulk of the stalled fork moved through the lesion (**Figure 2B**, Lanes 7-8). In comparison with the template without CPD in **Figure 2A**, this product is appreciably shorter than full length (e.g. compare **Figure 2B**, Lane 8 to **Figure 2A**, Lane 8). It is possible that Pol α becomes uncoupled from CMG during its relatively long TLS action at the lesion, subsequently blocked from further extension by template re-annealing in the wake of CMG unwinding. Interestingly, in **Figure 2B**, Lanes 9-10, Pol α-driven CPD bypass no longer occurs in the presence of Pol ε· Of note, CMGE exhibited normal replication up to the fork junction (**Figure 2B**, Lane 4), demonstrating an active Pol ε and competent leading strand replisome. We return to the effect of Pol ε on Pol α TLS activity further below.

### Pol α overexpression does not affect UV sensitivity in the absence of TLS Pols

Having observed and characterized robust CPD TLS activity from Pol α *in vitro*, we wanted to know whether it might be acting as a TLS *in vivo*. To query the effect of Pol α overexpression on TLS activity, we asked whether overexpression of Pol α could result in a measurable increase in DDT activity in response to UV exposure, relative to baseline. Importantly, this experiment is unable to inform us about the physiological relevance of Pol α TLS activity, nor can we devise one that would: unlike specialized TLS Pols, deletion of *POL1*, encoding the large polymerase subunit of Pol α, is known to be lethal in baker’s yeast [39], as is mutation of its polymerase active site in fission yeast [40], indicating that testing physiological TLS activity in this critical polymerase active site *in vivo* is likely not possible. Thus, we emphasize that the following experiment simply asks whether overexpression of Pol α can rescue cell viability in UV-perturbed TLS knockout strains, which could be relevant if high concentrations of Pol α were able to partially rescue UV sensitivity by way of either increased TLS activity or overcoming any potential suppression of Pol α TLS activity that may be taking place in the cell.

To eliminate activity from the known TLS Pols, we knocked out *REV1, REV3* (subunit of Pol ζ), and *RAD30* (encoding Pol η) to generate a ΔTLS strain. We further integrated *GAL1/10*-controlled overexpression cassettes of all four subunits of Pol α into this strain (Gal/Pol α). We observed by western blot robust expression of Pol1-FLAG after induction with galactose (Gal) and did not see any leaky expression in the absence of Gal (**Figure S2A**). Cells were grown in YP-Glycerol to avoid repressing Gal promoters with glucose and were exposed to 150 J/m^2^ UV-C light, with or without Gal induction. Consistent with other reports [37,38], we observe that UV exposure is well tolerated in the WT strain but is significantly more sensitive to UV in the ΔTLS strain; growing in YPG, less ΔTLS cells survive UV irradiation compared to WT cells (**Figure S2B**, top plates). The same effect can be seen when growing in both YPG and galactose (**Figure S2B**, bottom plates). Yet in all cases, the overexpression of Pol α did not appear to affect UV sensitivity (**Figure S2B**, compare UV sensitivity of strains bearing Gal/Pol α overexpression cassettes in the absence (top plates) or presence (bottom plates) of induction of Pol α overexpression by Gal). Thus, we conclude that increased nuclear concentrations of Pol α cannot increase its activity as a CPD TLS *in vivo*.

### Strong inhibition of Pol α TLS by Pol ε and RPA

Our observation that overexpression of Pol α could not rescue UV sensitivity in a ΔTLS background indicated that the intrinsic TLS activity of Pol <u>α</u> we report here may somehow be enzymatically inhibited in cells, even during replication stress. We saw above that we could not observe extension past the CPD lesion with 20 nM Pol ε present (compare **Figure 2B** lanes 9-10 to lanes 7-8), so we wanted to know whether this apparent inhibition could be competed with by increased concentrations of Pol α. Interestingly, 5 nM of Pol ε was able to inhibit CPD bypass by Pol α at the fork, which was not reversed when we titrated Pol α up to 80 nM. In all conditions where Pol ε is present, most of the product was stuck at the CPD lesion (**Figure 3A**), indicating that Pol ε strongly inhibits Pol α TLS activity. We further considered the possibility that Ctf4, a homotrimer that helps tether Pol α and other accessory factors to CMG helicase [39–42], may contribute to Pol α TLS activity. However, upon addition of Ctf4 to the reaction we saw similarly strong inhibition of Pol α by Pol ε regardless of Pol α concentration, indicating that Ctf4 likely does not coordinate the bypass activity of Pol α in a leading strand replisome (**Figure S3A**). We next asked whether the proofreading capability of Pol ε may be responsible for this suppression. Indeed, we saw that a 5’-3’ exonuclease deficient Pol ε (Pol ε^exo-^) was competent for replication, even exhibiting a very small amount of CPD lesion bypass alone after 30 minutes consistent with a shift towards polymerase activity and a high mutation rate [47,48], but could no longer suppress Pol α TLS activity (**Figure S4)**. We therefore consider it likely that exonuclease underlies the strong inhibition of Pol α TLS activity by Pol ε, but further studies are needed to confirm this. However, fast proofreading activity of a relatively slow incorporation across from a bulky lesion may explain why inhibition by Pol ε cannot be simply outcompeted by excess Pol α.

**Figure 3.**
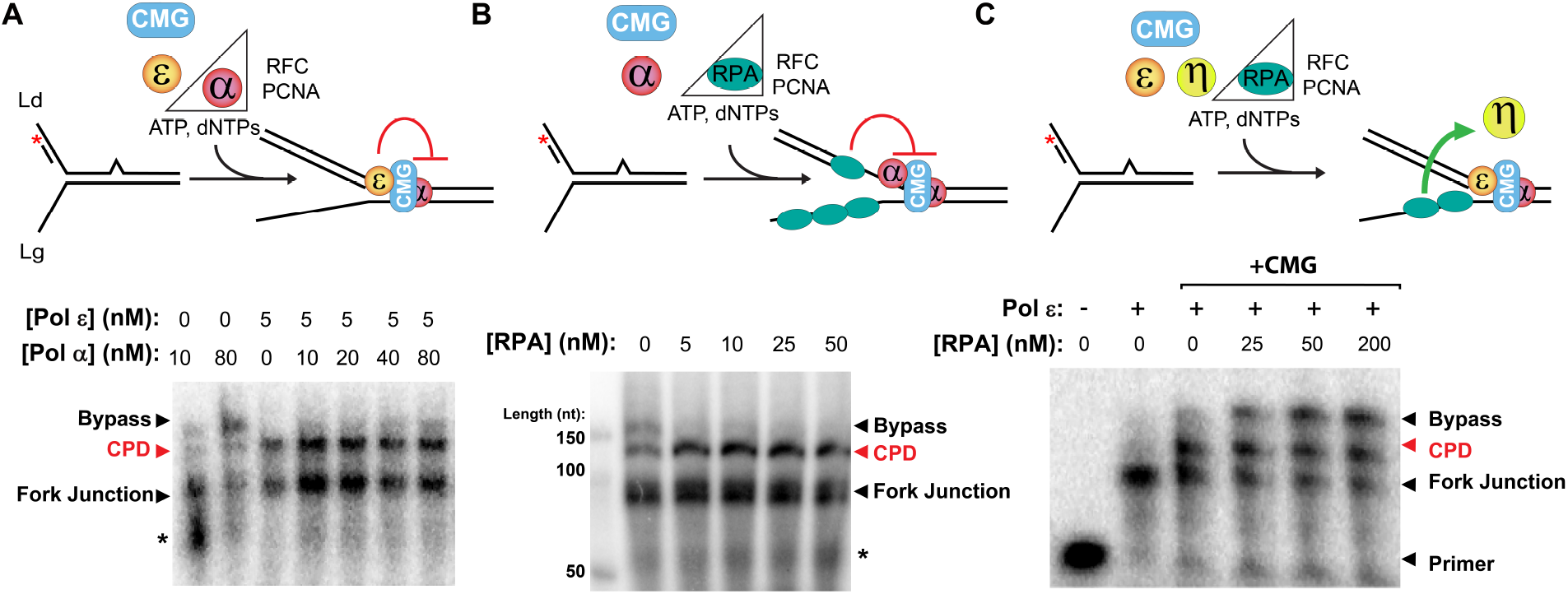
Pol ε and RPA strongly inhibit Pol α CPD TLS but stimulate Pol η CPD TLS. Primer extension assays showing **A)** A small amount of Pol ε inhibits Pol α CPD TLS activity and is not outcompeted by excess Pol α, **B)** Small amounts of RPA strongly inhibit Pol α TLS at the fork, and **C)** Pol η is not inhibited by Pol ε and is stimulated by RPA. Top (both panels): Reaction scheme indicating relevant additions. Bottom (both panels): Denaturing gel of primer extension assay. Reactions were performed with the indicated Pol and/or RPA concentrations, and all reactions were stopped at 20 min.

Next, we asked whether RPA could affect the TLS polymerase activity of Pol α, as it can often play a role in either stimulating or inhibiting replicative enzymes, e.g., it has been shown to inhibit primase activity of Pol α but does not inhibit and can even stimulate its polymerase activity [43,10]. Surprisingly, we saw that RPA strongly inhibits translesion activity of Pol α past a leading strand CPD (**Figure 3B)**. Interestingly, we observed that preincubation of a lagging strand template with increasing concentrations RPA did not appreciably weaken replication on undamaged templates in terms of percent full length product observed, but strongly inhibited Pol α TLS bypass efficiency (**Figure S5**). Having seen strong inhibition of Pol α TLS by Pol ε and RPA, we also asked whether RFC can inhibit leading strand TLS by Pol α, as we have shown it to robustly inhibit Pol ε, but not Pols δ or α, on the lagging strand [35]. However, we observed that RFC (either by competition or via clamp loading activity) did not affect bypass, and also that lack of clamp loading does not affect the reaction, consistent with the ability of both Pol ε and Pol α to work independently of PCNA sliding clamp (**Figure S5B)**. Further, we considered the possibility that Pol δ could inhibit Pol α TLS via nonspecific competition or through its polymerase activity. In order to probe the effect of Pol δ, the lagging strand polymerase, Pol α needs to make lagging strand primers, requiring NTPs, so this experiment also tests whether Pol α priming (either on the lagging strand or ahead of the junction) inhibits its TLS activity. However, in the presence or absence of RFC clamp loader, neither Pol δ or Pol α primase activity appeared to inhibit Pol α CPD TLS (**Figure S6**).

### Stimulation of Pol η CPD TLS by RPA and Pol ε

Pol η, the yeast TLS Pol that is specialized for pyrimidine dimers, has been shown to remodel a CPD-stalled fork for efficient bypass [50]. In that report, the presence of RPA is tolerated by Pol η, conversely to what we show here for Pol α TLS activity. Further, RPA has also been shown to stimulate the TLS activity of human Pol η [51]. Exploring this, we observed an apparent dose-dependent increase in activity of yeast Pol η with RPA addition (**Figure 3**). The fact that we observed Pol η CPD TLS activity in the presence of Pol ε, unlike what we observed for Pol α, indicates that Pol η is likely stimulated by lesion-stalled Pol ε; in fact, we have seen like others that Pol η on its own is weakly processive and instead relies on polymerase handoff for activity [52,53]. Taken together, these results demonstrate that RPA and Pol ε may be regulating the bypass of Pol α and Pol η through disparate mechanisms, potentially related to differences in lesion bypass rates and/or interactions between RPA or Pol ε and the respective TLS Pol.

### Pol α remains predominately in the open conformation to accommodate a CPD lesion

Having demonstrated the intrinsic capacity for bulky TLS by Pol α, we wanted to explore the mechanics of the ability of Pol α to read through and incorporate across a bulky lesion that is not known to be accommodated in replicative polymerase active sites. To monitor the distance between the thumb and fingers domains (a proxy for the width of the DNA binding cleft and polymerase active site), we constructed a version of Pol α with a LD555/LD655 (donor/acceptor) FRET pair, which we refer to as Pol α^555/655^, for measuring the FRET efficiency between the thumb and fingers domains during incorporation (schematic in **Figure 4A** and molecular model in **Figure S7**). We first tethered biotinylated primed ssDNA templates with or without a CPD (like Ld/P and Ld^CPD^/P used in Figure 1) onto a surface passivated flow cell and then added Pol α^555/655^ at a 1:5 ratio of protein:DNA to ensure DNA binding. Reaction conditions were identical to those in Figure 1, with the addition of a photoprotective oxygen scavenging system. We recorded movies immediately after flowing on all components to capture Pol α^555/655^ during nucleotide incorporation, under the assumption that most of the population of Pol α would be stalled at a CPD given the slow bypass kinetics we observe herein. We then quantified the apparent Fluorescence Resonance Energy Transfer efficiency (FRET E_app_) after semi-automatically filtering traces to remove spurious signals and post-photobleaching data (**Figure S8**). On an undamaged template/primer (T/P), we observed a broad FRET distribution (**Figure 4B**) that was largely composed of dynamic events like the representative traces shown in **Figure 4C**. This FRET trace highlights dynamic motion between the thumb and fingers domain, which is consistent with the conformational change of fingers clamping down on the dNTP and primer terminus in its palm domain that is associated with dNTP incorporation [35]; we thus refer to the high and low FRET states as “closed” and “open” states, respectively (**Figure 4A**).

**Figure 4.**
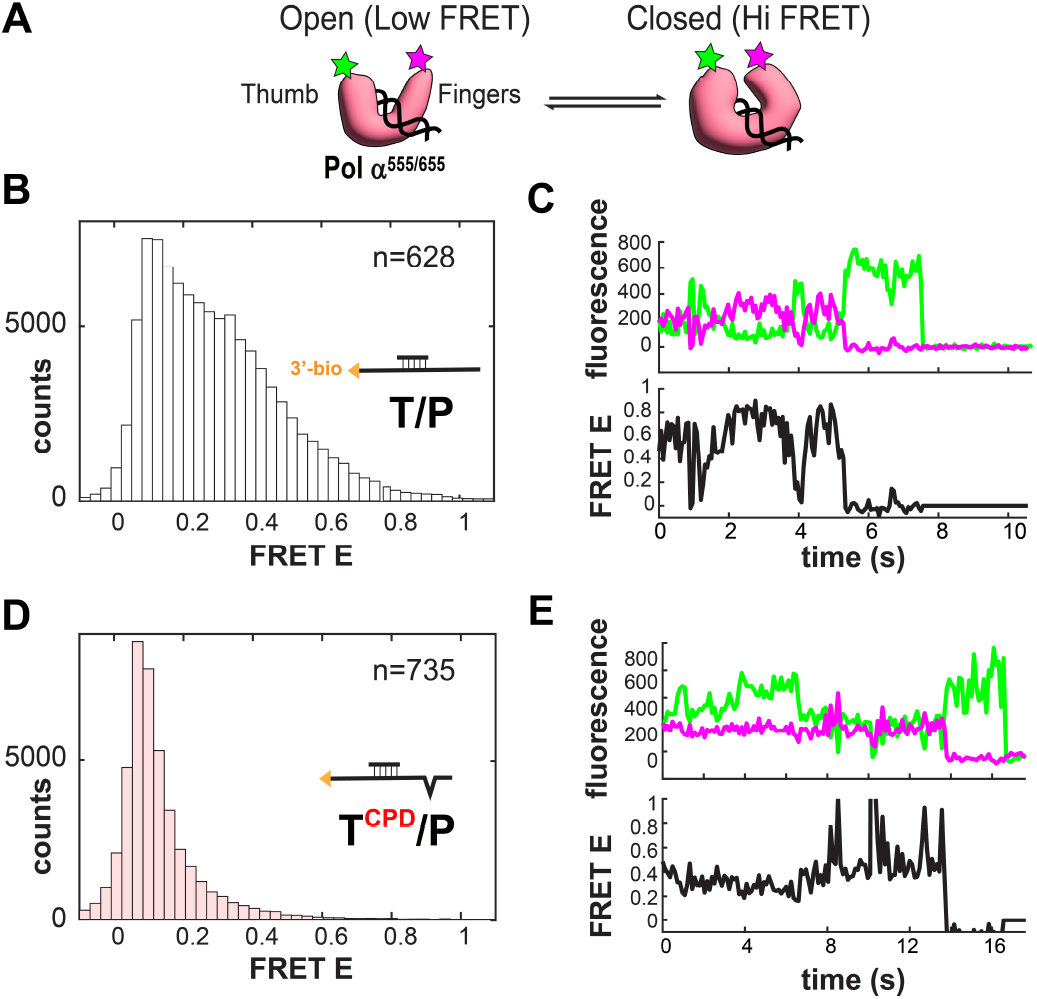
Pol α accommodates CPD lesions by shifting toward the open state. **A)** Schematic showing FRET construct. Pol α is labeled with the LD555 (green) and LD655 (red) FRET pair at the thumb and fingers domains. Imaging takes place during dNTP incorporation across from CPD on the primed template, which is identical to those used in Figure 1 but with 5’-biotin. With this experimental setup, large conformational changes consistent with opening and closing of the thumb and fingers domains around the DNA substrate can be measured. **B)** FRET distribution of thumb/fingers FRET during replication of an undamaged template (n=628 molecules). **C)** Representative trace showing FRET fluctuation with anticorrelated donor/acceptor signal as well as acceptor photobleaching followed by donor photobleaching. **D)** FRET distribution of thumb/fingers FRET during replication of the template/primer with a CPD (n=735). **E)** Representative FRET trace of an acceptor-then-donor photobleaching event.

Remarkably, when we flowed this same Pol α^555/655^ construct onto a T^CPD^/P substrate (i.e., CPD as the only variable change), we saw this distribution collapse into a single low FRET peak (**Figure 4D**). Although we saw a few dynamic events, there were significantly fewer of them, and they appeared to visit the “closed” state more seldomly (**Figure 4E**). Most of the traces, however, were relatively static low-FRET traces, indicating that Pol α largely stayed open while it was incorporating across from a CPD. In both **Figure 4** and **Figure S9**, we highlight traces with acceptor photobleaching followed by donor photobleaching as ideal FRET traces, largely because low FRET signals are difficult to distinguish between donor-only peaks (which we also eliminated during the filtering step). Further, by adding a donor:acceptor stoichiometry dimension to the histograms in Figure 3, we show in **Figure S10** that the data fell within the acceptable range of E_app_ between ∼0.25 and ∼0.75 and centered on ∼0.5, indicating relatively equal donor:acceptor intensity stoichiometry in the filtered traces. Collectively, the data indicate that Pol α likely stays largely open during bypass of a bulky CPD. Further structural evidence is required to corroborate this result. Still, our observation that CPD TLS requires Pol α to visit its relatively inaccessible open conformation may help explain the slow bypass kinetics that we observe.

## DISCUSSION

In this report, we demonstrate that Pol α possesses CPD TLS activity *in vitro*. To our knowledge, this is the first report of a replicative polymerase incorporating across a bulky lesion. Other groups have reported TLS activity in replicative polymerases. Interestingly, one report demonstrated desensitization to 4-NQO damage (which results in 8-oxoG lesions) upon increasing dNTP production in a yeast strain in which all TLS enzymes were removed, suggesting that at least one replicative Pol might be able to bypass CPDs; this study also found that Pol ε could bypass 8-oxoG lesions when increasing dNTP concentration [45]. Pol δ has been demonstrated to bypass 8-oxoG lesions, albeit inefficiently compared to Pol η in humans [46]. It has observed that Pol δ and Pol ε coordinate replicase switching to bypass thymine glycol and 8-oxo-G lesions in yeast [47]. However, these Pols could not bypass CPDs, consistent with a previous report demonstrating that Pol δ or ε may be involved in TLS proofreading via their exonuclease activities but could not incorporate nucleotides opposite a CPD [30]. Indeed, Pol ε appears to retain limited TLS activity, but only for non-bulky lesions like abasic sites [45,48], and, as we show here, can be pushed toward very moderate CPD bypass by removing its proofreading activity. Two studies have demonstrated bypass of CPD lesions by Pol δ at dNTP levels of 0.1 mM or greater [20,21], which, like exonuclease mutations, can cause mutator phenotypes by shifting the kinetics towards polymerization. Hence, to the best of our knowledge, replicative polymerases at physiological dNTP levels have not been shown to bypass sterically bulky lesions. Here, we show that Pol α can polymerize through a CPD either on the leading or lagging strand. We provide evidence that its DNA binding groove likely remains open while accommodating a CPD relative to an undamaged template, further indicating that the Pol α active site has intrinsic steric capacity competent to embrace a bulky lesion during synthesis. We also show that RPA and Pol ε each strongly inhibit Pol α CPD TLS by Pol α. In the case of TLS inhibition by Pol ε, this appears to be independent of relative polymerase stoichiometry, possibly owing to suppression of Pol α TLS activity by proofreading. Consistently, we observed that Pol α overexpression cannot overcome UV sensitivity in a ΔTLS background. We interpret this to mean that inherent Pol α TLS is suppressed by the replisome itself during normal, unchallenged replication. Physiologically, this tracks with the idea that specialized error-prone TLS Pols evolved to be specifically, rapidly, and—critically—temporarily recruited to remodel lesion-stalled replisomes and not on unchallenged elongating forks.

It was previously shown that repriming past a CPD lesion is efficient on the lagging strand but inefficient on the leading strand [10], a result that we have also observed when probing for sequence-specific incorporation of ^32^P-α-NTPs past a CPD lesion in a synthetic nucleotide-biased template. In our hands, repriming past a CPD was undetectable on leading strand templates working with a reconstituted replisome (not shown). Although it was demonstrated that primase activity is responsible for most of CPD bypass on the leading strand, we consider the possibility that the TLS activity of Pol α that we report here could be at least partially responsible for bypassing leading strand CPD lesions in that and other recent studies of lesion skipping at the replisome [10,29]. Since replication is initiated at origins in those studies, requiring primase activity of Pol α, NTPs and/or Pol ε cannot be omitted, making it difficult in some instances to discern between repriming past the CPD lesion on the leading strand and potential CPD TLS. That report also demonstrated that depletion of RPA-stimulated lesion skipping by repriming but showed that RPA did not inhibit polymerase activity [10]. In agreement with this, we saw that RPA did not inhibit replication on undamaged templates, but that it did strongly inhibit TLS. Thus, RPA may somehow only inhibit Pol α translesion activity, perhaps exploiting its relatively slow bypass kinetics. Relatedly, though we observe that Pol α works with CMG helicase to bypass a leading strand CPD, recent cryoEM structures revealed that Pol α preferentially binds CMG on the lagging strand side, restricting access to the leading strand [27], indicating the possibility that Pol α is working distributively and nonspecifically behind CMG in its capacity as a TLS Pol. We showed that Pol ε, which tightly binds CMG [34,35], was strongly inhibitory to Pol α TLS, even at small amounts. Even though Pol ε has been shown to readily exchange in an active fork [50], this inhibition could not be outcompeted by relatively higher Pol α concentration. It has been clearly demonstrated that, upon encountering a CPD, CMG uncouples from Pol ε and continues to unwind [10], thus, outside of any inhibition of TLS activity that may arise due to proofreading by Pol ε, we consider it likely that the relative affinity of Pol α for engaging a CPD lesion is much lower than that of Pol ε, which may remain stably engaged at a lesion until rescued by a DDT pathway. More work is required to fully understand the nature of Pol α activity at stalled replication forks.

It remains unclear whether Pol α has any role at replication forks *in vivo* or whether this activity has played a role in damage tolerance throughout evolutionary history*—*it is entirely possible that early versions of Pol α and/or Pol1 played a role as a TLS Pol in ancestral organisms. Further, since RPA strongly inhibits Pol α TLS in this report and was also reported to inhibit lesion skipping [10], it seems possible that the role of Pol α in negotiating a stalled replication fork (whether via repriming or translesion synthesis) has been relegated to rarely invoked yet vital tasks through negative regulation. For instance, cells may have evolved to invoke Pol α TLS strictly during “replication catastrophe,” critical instances of replication stress where RPA is fully exhausted, leaving the genome vulnerable to deleterious nucleolytic attack [60,61]. It is also possible that TLS via Pol α is error prone compared to canonical TLS Pols such that replisome architecture evolved to constitutively inhibit its activity at the fork entirely. Future studies are required to better understand these processes.

## METHODS

### Strains and plasmids

The base strain used for constructing overexpression strains is OY01 (W303; *ade2-1 ura3-1 his3-11,15 trp1-1 leu2-3,112 can1-100 bar1Δ MATa pep4::KANMX6*) [53]. Relevant plasmids and DNA are listed in Table S1.

### Protein purification

The purification of Pol α, Pol δ, Pol ε, Pol ε^exo-^ (POL2^D290A/E292A^), CMG, Mcm10, RFC, PCNA, RPA, Ctf4, and Sfp has been previously described [33,38,62,63]. The purity of protein complexes was assessed by Coomassie stained SDS-PAGE, and all enzymes were assessed for activity *in vitro* before use in assays.

### Pol α^555/655^ purification

A synthetic Pol1 fragment (Twist Bioscience; Table S1) containing small (12 AA) peptides called S6—which *B*.*subtilis* 4’-phosphopantetheinyl transferase (Sfp) can covalently attach CoA-conjugated fluorophores to with high efficiency [64,65]— was used to replace the thumb and finger region of the *3xFLAG*_ *POL1_ pRS402_GALt* integration vector using Gibson assembly (NEB), after digesting the vector with BmgBI and PstI. The S6 in the finger domain was inserted between S959 & R960; the thumb S6 was inserted between K1138 and A1139 (illustrated in Figure S3.1). The resultant *3xFLAG_Pol1*^*S6/S6*^ construct was integrated into a strain containing *GAL*/*Pri2, GAL/Pol12*, and *GAL/Pri1*. The donor also contained silent AflII sites in the fingers/thumb regions to facilitate colony selection. Next, the 4-subunit protein complex was overexpressed and purified on anti-FLAG resin as previously described for Pol α[38]. The complex was then labeled with the 555/655 FRET pair using 1:2:5:5 ratio of Pol α^S6/S6^, Sfp, and LD555-CoA, and LD6555-CoA (Lumidyne Technologies), and labeling for 2 hrs at 25°C in 50 mM Tris-HCl, pH 6.8, 5% glycerol, 150 mM NaCl, 10 mM MgCl2, and 1mM DTT. Total labeling efficiency of both sites was >90%, assessed by absorbance at 555 and 655 vs protein concentration obtained by Bradford assay. Sfp and excess dye were removed with a Sephadex 200 gel filtration column (GE healthcare) with a mobile phase of 50 mM Tris-HCl, pH 7.5, 2% glycerol, 150 mM NaCl, and 1mM DTT. Peak fractions were flash frozen, aliquoted, and stored at -80°C.

### Pol η purification

Pol η was made by cloning and integrating *3xFLAG-Rad30* into OY01 via *pRS403*-*GALt* (HIS3::GAL integration vector with terminator[66]; Table S1). Cells were grown under auxotrophic (-His) selection 30°C in Yeast Peptone (YP)-glucose, split into 12L YP-glycerol, grown to an OD_600_ of 0.6 at 30°C, and induced with 2% galactose for 6 hours. Cells were harvested by centrifugation at 1,500 x g, resuspended with Buffer A (25 mM HEPES pH 7.5, 0.5mM TCEP with 10% glyercol and 400 mM NaCl) and protease inhibitors (Sigma P8215), frozen dropwise into liquid nitrogen, and lysed with a cryogenic grinding mill at for 12 cycles of 5 min cool followed by 2 min grind at 9 cycles per second (SPEX SamplePrep). The lysate was diluted with 20 mL Buffer A and clarified by centrifugation at 43,500 x g in an JA-20 rotor for 2 h at 4°C. The supernatant was applied to a 1 mL anti-FLAG M2 column (Sigma) and washed with 30 column volumes (CV) buffer A. Protein was eluted in 5 CV Buffer A including 0.3 mg/mL 3xFLAG peptide (EZBiolab). The eluate was loaded onto a 1 mL Mono-Q column (GE healthcare) and eluted with a 10 CV 0.1–1.5 M NaCl gradient (in Buffer A base, with 5% glycerol). Peak fractions were flash frozen, aliquoted, and stored at -80°C.

### Primer extension templates

Template construction has been previously described, which involves ligating two oligos together and purifing the 243 nt leading strand [59] (see Table S1 for oligos). A 30 nt primer is annealed to this template to make ssDNA substrates with (Ld^CPD^/P) or without (Ld/P) CPD. To make replication forks, these constructs were annealed with a 185 nt lagging strand, providing a noncomplementary fork for CMG loading and a 125 nt duplex. Synthetic oligonucleotides containing CPD were ordered from Trilink Biotechnologies; all other were oligos were from IDT. Primer 5’-^32^P labeling was with PNK (NEB) and ^32^P-γ-ATP (Revvity) according to manufacturer protocol.

### Primer extension assays

Reaction volumes were 25 μL and contained 0.5 nM of the template, primed with a 5’-^32^P-primer (see Table S1 for sequences). Reactions contained 25 mM Tris-OAc pH 7.5, 5% (vol/vol) glycerol, 50 μg/mL BSA, 2 mM TCEP, 3 mM DTT, 10 mM Mg-OAc, 50 mM potassium glutamate, mM EDTA, and all stages were carried out at 30°C. For primed ssDNA reactions, replication was initiated by incubating the template with 20 nM Pol α and 60 μM dNTP unless otherwise noted. For forked templates, DNA was first loaded with 20 nM CMG and 40 nM Mcm10 for 10 min in the presence of 0.1 mM ATP—sufficient for CMG loading on the fork but not dsDNA unwinding [33]. Unless otherwise indicated, polymerases were introduced at 20 nM, with 60 nM PCNA and 5 nM RFC for 5 min. Where relevant (e.g., for RPA and Pol ε titrations) the salt from added proteins was balanced so all reactions were isotonic. Replication was initiated by the addition of 5 mM ATP and 60 μM dNTPs, and were stopped at the indicated times with a STOP solution containing 1% SDS and 40 mM EDTA. Reactions were run on 10% denaturing PAGE (8M Urea) at 25 mA for 40 min. Gels were backed with Whatman 3MM filter paper, exposed to a storage phosphor screen for 16 hours, and visualized on a Typhoon FLA 9500 PhosphorImager.

### UV sensitivity assay

TLS knockout (ΔTLS) strains were created by knocking out and replacing *REV1, REV3* (a subunit of Pol ζ) and *RAD30* (Pol η) with nutritional (LEU2, TRP) and antibiotic (NTC) markers in *Saccharomyces cerevisiae* strains OY01 (described above) and OY03, a Pol α overexpression strain containing integrated copies of *GAL-pol1*-*3xFLAG, GAL*-*Pri2, GAL-Pol12*, and *GAL-Pri1*)[38]. All growths were performed at 30°C with shaking at 220rpm. Sequence verified single colonies were inoculated into 5 mL YPD and grown for 16 h, diluted to OD_600_= 0.2 in 5 mL YPG (2% glycerol) and grown for 7–8 h. Cultures were diluted into 50 mL YPG (OD_600_= 0.008) and grown for 16 h. Cultures (OD_600_∼0.4–0.6) were either grown in YPG or induced with 2% galactose for 3 h. Cells (5 OD_600_ units) were then harvested, washed, and resuspended in 5 mL of either 2% glycerol (uninduced) or 2% galactose (induced). Cell suspensions were transferred to Petri dishes and, where indicated, irradiated with UV-C (254 nm) at 150 J/m^2^ (∼0.1 mW/cm^2^ for 2.5 min). ΔTLS and wild-type strains were then assessed for UV sensitivity after treatment using a serial dilution spot assay in the presence or absence of 2% galactose (5000, 500, 50, 5 cells/µL; 4 µL of each dilution was spotted onto YPG plates).

### Western Blot

Cells lysis was performed as described [71]. 2.5 OD_600_ units of yeast were pelleted 5 min at 1000xg and resuspended in 100 mL of water. 0.1 mL of 0.2 M NaOH was added for 5 min at 25°C and spun for 30 sec at 5000xg. The pellet was resuspended in 60 µl of 2X sample prep buffer (250 mM Tris pH 6.8, 10% glycerol, 1% SDS, 10% β-mercaptoethanol, 0.005% bromophenol blue), boiled 3 min, and centrifuged 1 min at 21,000xg, collecting supernatant. 8 µl per lane was loaded on a 7.5% acrylamide gel for SDS-PAGE and transferred to nitrocellulose for western blotting. Membrane was blocked in 1% BSA in TBS (10 mM Tris pH 8.0, 150 mM NaCl). Antibody incubations and washes were in TBST (10 mM Tris pH 8.0, 150 mM NaCl, 0.1% Tween-20) with 1% BSA. Primary antibodies: mouse anti-FLAG (Sigma-Aldrich #F18041) at 1:2000, or mouse anti-α-tubulin (ABM # G094) at 1:8000 for a loading control. Secondary antibodies: DyLight goat anti-mouse 680 nm. Nitrocellulose membrane was imaged on a LI-COR Odyssey Clx imaging system.

### Single-molecule FRET assays

Single-molecule FRET experiments were performed on a homebuilt prism-based total internal reflection (TIRF) setup based on a Nikon Ti2-E inverted fluorescence microscope. Flow cells were constructed from Quartz slides (G. Finkenbeiner, Inc.) and an ∼80 uM adhesive gasket (ARcare 90445). Slides were PEGylated with mPEG and bioPEG (Laysan Bio) and incubated with Neutravidin (Sigma) as described[67]. Biotin-conjugated DNA was surface-tethered at 250 pM and Pol α^555/655^ was incubated at 50 pM. General assay conditions were identical to those used in the primer extension assays, including 60 μM dNTPs, and additionally included the protocatechuic acid/protocatechuate-3,4-dioxygenase (PCA/PCD) oxygen scavenging system[68] with 40 nM PCD, 2.5 mM PCA, 1 mM Trolox, 1 mM cyclooctatetraene, and 1 mM 4-nitrobenzyl alcohol to prevent photobleaching and/or blinking. For each experimental repetition (n=6) performed to validate the observation, the tested conditions (i.e., undamaged vs damaged template) were recorded on two channels of the same slide. Continuous wave lasers (488 nm/100 mW, 532 nm/80 mW, and 640 nm/75 mW; Coherent OBIS) were used continuously or alternately by exciting 555 and 655 dyes every other frame, a.k.a. Alternating Laser Excitation (ALEX). Emission was collected with a 60X Plan Apo TIRF water immersion objective, sent through an image splitter (Cairn) with a 560 nM longpass (LP) dichroic mirror (Chroma T560lpxr-UF1) and a 640 nM LP dichroic (Chroma T640lpxr-UF2). The 488 and 555 dye emissions were filtered through cleanup filters (Chroma ET525/50m and 585/65m, respectively). Spectrally filtered images were projected onto the focal plane of three scientific complementary metal-oxide Semiconductor (sCMOS) cameras (ORCA Fusion, Hamamatsu), and movies were recorded at a rate of 86 ms per frame. The data was parsed using custom Python scripts and analyzed using SPARTAN[69] and iSMS[70]. All FRET trajectories were filtered to remove post-photobleaching data, and ALEX was used to select traces with FRET pairs to eliminate choosing donor-only peaks.

## ACKNOWLEDGEMENTS

We thank the O’Donnell lab at Rockefeller University for sharing strains and integration vectors.

## FUNDING SOURCES

Funding was provided by NIGMS (R00GM126143 and R35GM147105 to G.D.S.).

The Khorana Program for Scholars (Department of Biotechnology, India, Indo-U.S. Science and Technology Forum, and WINStep Forward) supported A.S.

## REFERENCES

[1] D.J. Chang, K.A. Cimprich, DNA damage tolerance: when it’s OK to make mistakes, Nat Chem Biol 5 (2009) 82–90. 10.1038/nchembio.139.

[2] S.P. Bell, K. Labib, Chromosome Duplication in Saccharomyces cerevisiae, Genetics 203 (2016) 1027–1067. 10.1534/genetics.115.186452.

[3] P.M.J. Burgers, T.A. Kunkel, Eukaryotic DNA Replication Fork, Annual Review of Biochemistry 86 (2017) 417–438. 10.1146/annurev-biochem-061516-044709.

[4] M.F. Goodman, R. Woodgate, Translesion DNA Polymerases, Cold Spring Harb Perspect Biol 5 (2013). 10.1101/cshperspect.a010363.

[5] K.J. Marians, Lesion Bypass and the Reactivation of Stalled Replication Forks, Annual Review of Biochemistry 87 (2018) null. 10.1146/annurev-biochem-062917-011921.

[6] K.T. Powers, M.T. Washington, Eukaryotic translesion synthesis: Choosing the right tool for the job, DNA Repair 71 (2018) 127–134. 10.1016/j.dnarep.2018.08.016.

[7] W. Yang, Y. Gao, Translesion and Repair DNA Polymerases: Diverse Structure and Mechanism, Annu Rev Biochem 87 (2018) 239–261. 10.1146/annurev-biochem-062917-012405.

[8] T.A. Guilliam, J.T.P. Yeeles, Reconstitution of translesion synthesis reveals a mechanism of eukaryotic DNA replication restart, Nat Struct Mol Biol 27 (2020) 450–460. 10.1038/s41594-020-0418-4.

[9] J.D. Kaszubowski, M.A. Trakselis, Beyond the Lesion: Back to High Fidelity DNA Synthesis, Front. Mol. Biosci. 8 (2022). 10.3389/fmolb.2021.811540.

[10] M.R.G. Taylor, J.T.P. Yeeles, The Initial Response of a Eukaryotic Replisome to DNA Damage, Molecular Cell 70 (2018) 1067-1080.e12. 10.1016/j.molcel.2018.04.022.

[11] R.E. Johnson, S. Prakash, L. Prakash, Efficient Bypass of a Thymine-Thymine Dimer by Yeast DNA Polymerase, Polη, Science 283 (1999) 1001–1004. 10.1126/science.283.5404.1001.

[12] A. Vaisman, R. Woodgate, Translesion DNA polymerases in eukaryotes: what makes them tick?, Crit Rev Biochem Mol Biol 52 (2017) 274–303. 10.1080/10409238.2017.1291576.

[13] J. Trincao, R.E. Johnson, C.R. Escalante, S. Prakash, L. Prakash, A.K. Aggarwal, Structure of the catalytic core of S. cerevisiae DNA polymerase eta: implications for translesion DNA synthesis, Mol Cell 8 (2001) 417–426. 10.1016/s1097-2765(01)00306-9.

[14] C. Biertümpfel, Y. Zhao, Y. Kondo, S. Ramón-Maiques, M. Gregory, J.Y. Lee, C. Masutani, A.R. Lehmann, F. Hanaoka, W. Yang, Structure and mechanism of human DNA polymerase η, Nature 465 (2010) 1044–1048. 10.1038/nature09196.

[15] Z. Zhuang, R.E. Johnson, L. Haracska, L. Prakash, S. Prakash, S.J. Benkovic, Regulation of polymerase exchange between Polη and Polδby monoubiquitination of PCNA and the movement of DNA polymerase holoenzyme, Proceedings of the National Academy of Sciences 105 (2008) 5361–5366. 10.1073/pnas.0801310105.

[16] K. Yang, C.P. Weinacht, Z. Zhuang, Regulatory Role of Ubiquitin in Eukaryotic DNA Translesion Synthesis, Biochemistry 52 (2013) 3217–3228. 10.1021/bi400194r.

[17] B.D. Freudenthal, L. Gakhar, S. Ramaswamy, M.T. Washington, Structure of monoubiquitinated PCNA and implications for translesion synthesis and DNA polymerase exchange, Nat Struct Mol Biol 17 (2010) 479–484. 10.1038/nsmb.1776.

[18] A. Bernad, A. Zaballos, M. Salas, L. Blanco, Structural and functional relationships between prokaryotic and eukaryotic DNA polymerases., EMBO J 6 (1987) 4219–4225.

[19] M. O’Donnell, L. Langston, B. Stillman, Principles and Concepts of DNA Replication in Bacteria, Archaea, and Eukarya, Cold Spring Harb Perspect Biol 5 (2013) a010108. 10.1101/cshperspect.a010108.

[20] C.L. O’Day, P.M. Burgers, J.S. Taylor, PCNA-induced DNA synthesis past cis-syn and trans-syn-I thymine dimers by calf thymus DNA polymerase delta in vitro, Nucleic Acids Res 20 (1992) 5403–5406. 10.1093/nar/20.20.5403.

[21] T. Narita, T. Tsurimoto, J. Yamamoto, K. Nishihara, K. Ogawa, E. Ohashi, T. Evans, S. Iwai, S. Takeda, K. Hirota, Human replicative DNA polymerase δcan bypass T-T (6-4) ultraviolet photoproducts on template strands, Genes Cells 15 (2010) 1228–1239. 10.1111/j.1365-2443.2010.01457.x.

[22] S. García-Gómez, A. Reyes, M.I. Martínez-Jiménez, E.S. Chocrón, S. Mourón, G. Terrados, C. Powell, E. Salido, J. Méndez, I.J. Holt, L. Blanco, PrimPol, an Archaic Primase/Polymerase Operating in Human Cells, Molecular Cell 52 (2013) 541–553. 10.1016/j.molcel.2013.09.025.

[23] J. Bianchi, S.G. Rudd, S.K. Jozwiakowski, L.J. Bailey, V. Soura, E. Taylor, I. Stevanovic, A.J. Green, T.H. Stracker, H.D. Lindsay, A.J. Doherty, PrimPol Bypasses UV Photoproducts during Eukaryotic Chromosomal DNA Replication, Mol Cell 52 (2013) 566–573. 10.1016/j.molcel.2013.10.035.

[24] E.O. Boldinova, A.V. Yudkina, E.S. Shilkin, D.I. Gagarinskaya, A.G. Baranovskiy, T.H. Tahirov, D.O. Zharkov, A.V. Makarova, Translesion activity of PrimPol on DNA with cisplatin and DNA–protein cross-links, Sci Rep 11 (2021) 17588. 10.1038/s41598-021-96692-y.

[25] S. Mourón, S. Rodriguez-Acebes, M.I. Martínez-Jiménez, S. García-Gómez, S. Chocrón, L. Blanco, J. Méndez, Repriming of DNA synthesis at stalled replication forks by human PrimPol, Nature Structural & Molecular Biology 20 (2013) 1383–1389. 10.1038/nsmb.2719.

[26] D. González-Acosta, E. Blanco-Romero, P. Ubieto-Capella, K. Mutreja, S. Míguez, S. Llanos, F. García, J. Muñoz, L. Blanco, M. Lopes, J. Méndez, PrimPol-mediated repriming facilitates replication traverse of DNA interstrand crosslinks, EMBO J 40 (2021) e106355. 10.15252/embj.2020106355.

[27] R. Mayle, R. Georgescu, M.E. O’Donnell, DNA polymerase α-primase can function as a translesion DNA polymerase, Proceedings of the National Academy of Sciences 122 (2025) e2517556122. 10.1073/pnas.2517556122.

[28] Z. Yuan, R. Georgescu, H. Li, M.E. O’Donnell, Molecular choreography of primer synthesis by the eukaryotic Pol α-primase, Nat Commun 14 (2023) 3697. 10.1038/s41467-023-39441-1.

[29] M.L. Jones, V. Aria, Y. Baris, J.T.P. Yeeles, How Pol α-primase is targeted to replisomes to prime eukaryotic DNA replication, Mol Cell 83 (2023) 2911-2924.e16. 10.1016/j.molcel.2023.06.035.

[30] E.A. Mullins, L.E. Salay, C.L. Durie, N.P. Bradley, J.E. Jackman, M.D. Ohi, W.J. Chazin, B.F. Eichman, A mechanistic model of primer synthesis from catalytic structures of DNA polymerase α-primase, Nat Struct Mol Biol 31 (2024) 777–790. 10.1038/s41594-024-01227-4.

[31] S.A.N. McElhinny, D. Kumar, A.B. Clark, D.L. Watt, B.E. Watts, E.-B. Lundström, E. Johansson, A. Chabes, T.A. Kunkel, Genome instability due to ribonucleotide incorporation into DNA, Nat Chem Biol 6 (2010) 774–781. 10.1038/nchembio.424.

[32] S.A. Nick McElhinny, B.E. Watts, D. Kumar, D.L. Watt, E.-B. Lundström, P.M.J. Burgers, E. Johansson, A. Chabes, T.A. Kunkel, Abundant ribonucleotide incorporation into DNA by yeast replicative polymerases, Proceedings of the National Academy of Sciences 107 (2010) 4949–4954. 10.1073/pnas.0914857107.

[33] G. Schauer, J. Finkelstein, M. O’Donnell, In vitro Assays for Eukaryotic Leading/Lagging Strand DNA Replication, Bio Protoc 7 (2017). 10.21769/BioProtoc.2548.

[34] L.D. Langston, D. Zhang, O. Yurieva, R.E. Georgescu, J. Finkelstein, N.Y. Yao, C. Indiani, M.E. O’Donnell, CMG helicase and DNA polymerase form a functional 15-subunit holoenzyme for eukaryotic leading-strand DNA replication, Proceedings of the National Academy of Sciences 111 (2014) 15390–15395. 10.1073/pnas.1418334111.

[35] G.D. Schauer, M.E. O’Donnell, Quality control mechanisms exclude incorrect polymerases from the eukaryotic replication fork, Proceedings of the National Academy of Sciences 114 (2017) 675–680. 10.1073/pnas.1619748114.

[36] M.R.G. Taylor, J.T.P. Yeeles, Dynamics of Replication Fork Progression Following Helicase–Polymerase Uncoupling in Eukaryotes, Journal of Molecular Biology 431 (2019) 2040–2049. 10.1016/j.jmb.2019.03.011.

[37] S.D. McCulloch, R.J. Kokoska, O. Chilkova, C.M. Welch, E. Johansson, P.M.J. Burgers, T.A. Kunkel, Enzymatic switching for efficient and accurate translesion DNA replication, Nucleic Acids Research 32 (2004) 4665–4675. 10.1093/nar/gkh777.

[38] R.E. Georgescu, G.D. Schauer, N.Y. Yao, L.D. Langston, O. Yurieva, D. Zhang, J. Finkelstein, M.E. O’Donnell, Reconstitution of a eukaryotic replisome reveals suppression mechanisms that define leading/lagging strand operation, eLife 4 (2015) e04988. 10.7554/eLife.04988.

[39] M.J. Carson, L. Hartwell, CDC17: an essential gene that prevents telomere elongation in yeast, Cell 42 (1985) 249–257. 10.1016/s0092-8674(85)80120-3.

[40] D. Bhaumik, T.S. Wang, Mutational effect of fission yeast pol alpha on cell cycle events, Mol Biol Cell 9 (1998) 2107–2123. 10.1091/mbc.9.8.2107.

[41] A.L. Abdulovic, S. Jinks-Robertson, The in Vivo Characterization of Translesion Synthesis Across UV-Induced Lesions in Saccharomyces cerevisiae: Insights Into Polζ- and Polη-Dependent Frameshift Mutagenesis, Genetics 172 (2006) 1487–1498. 10.1534/genetics.105.052480.

[42] J.P. McDonald, A.S. Levine, R. Woodgate, The Saccharomyces cerevisiae RAD30 Gene, a Homologue of Escherichia coli dinB and umuC, Is DNA Damage Inducible and Functions in a Novel Error-Free Postreplication Repair Mechanism, Genetics 147 (1997) 1557–1568. 10.1093/genetics/147.4.1557.

[43] A. Gambus, F. van Deursen, D. Polychronopoulos, M. Foltman, R.C. Jones, R.D. Edmondson, A. Calzada, K. Labib, A key role for Ctf4 in coupling the MCM2-7 helicase to DNA polymerase α within the eukaryotic replisome, The EMBO Journal 28 (2009) 2992– 3004. 10.1038/emboj.2009.226.

[44] A.C. Simon, J.C. Zhou, R.L. Perera, F. van Deursen, C. Evrin, M.E. Ivanova, M.L. Kilkenny, L. Renault, S. Kjaer, D. Matak-Vinkovic, K. Labib, A. Costa, L. Pellegrini, A Ctf4 trimer couples the CMG helicase to DNA polymerase [agr] in the eukaryotic replisome, Nature 510 (2014) 293–297.

[45] Z. Yuan, R. Georgescu, R. de L.A. Santos, D. Zhang, L. Bai, N.Y. Yao, G. Zhao, M.E. O’Donnell, H. Li, Ctf4 organizes sister replisomes and Pol α into a replication factory, eLife 8 (n.d.) e47405. 10.7554/eLife.47405.

[46] F. Villa, A.C. Simon, M.A. Ortiz Bazan, M.L. Kilkenny, D. Wirthensohn, M. Wightman, D. Matak-Vinkovíc, L. Pellegrini, K. Labib, Ctf4 Is a Hub in the Eukaryotic Replisome that Links Multiple CIP-Box Proteins to the CMG Helicase, Mol Cell 63 (2016) 385–396. 10.1016/j.molcel.2016.06.009.

[47] Y. Kawasaki, A. Sugino, Yeast Replicative DNA Polymerases and Their Role at the Replication Fork, Molecules and Cells 12 (2001) 277–285. 10.1016/S1016-8478(23)25248-6.

[48] J.A. St Charles, S.E. Liberti, J.S. Williams, S.A. Lujan, T.A. Kunkel, Quantifying the contributions of base selectivity, proofreading and mismatch repair to nuclear DNA replication in Saccharomyces cerevisiae, DNA Repair (Amst) 31 (2015) 41–51. 10.1016/j.dnarep.2015.04.006.

[49] K.A. Braun, Y. Lao, Z. He, C.J. Ingles, M.S. Wold, Role of Protein−Protein Interactions in the Function of Replication Protein A (RPA): RPA Modulates the Activity of DNA Polymerase α by Multiple Mechanisms, Biochemistry 36 (1997) 8443–8454. 10.1021/bi970473r.

[50] T.A. Guilliam, J.T.P. Yeeles, Reconstitution of translesion synthesis reveals a mechanism of eukaryotic DNA replication restart, Nat Struct Mol Biol 27 (2020) 450–460. 10.1038/s41594-020-0418-4.

[51] L. Haracska, R.E. Johnson, I. Unk, B. Phillips, J. Hurwitz, L. Prakash, S. Prakash, Physical and functional interactions of human DNA polymerase eta with PCNA, Mol Cell Biol 21 (2001) 7199–7206. 10.1128/MCB.21.21.7199-7206.2001.

[52] M.T. Washington, R.E. Johnson, S. Prakash, L. Prakash, Fidelity and Processivity of Saccharomyces cerevisiae DNA Polymerase η *, Journal of Biological Chemistry 274 (1999) 36835–36838. 10.1074/jbc.274.52.36835.

[53] A. Bresson, R.P.P. Fuchs, Lesion bypass in yeast cells: Pol η participates in a multi-DNA polymerase process, EMBO J 21 (2002) 3881–3887. 10.1093/emboj/cdf363.

[54] K.A. Johnson, Conformational coupling in DNA polymerase fidelity, Annu Rev Biochem 62 (1993) 685–713. 10.1146/annurev.bi.62.070193.003345.

[55] N. Sabouri, J. Viberg, D.K. Goyal, E. Johansson, A. Chabes, Evidence for lesion bypass by yeast replicative DNA polymerases during DNA damage, Nucl. Acids Res. 36 (2008) 5660– 5667. 10.1093/nar/gkn555.

[56] S.D. McCulloch, R.J. Kokoska, P. Garg, P.M. Burgers, T.A. Kunkel, The efficiency and fidelity of 8-oxo-guanine bypass by DNA polymerases delta and eta, Nucleic Acids Res 37 (2009) 2830–2840. 10.1093/nar/gkp103.

[57] T.A. Guilliam, J.T. Yeeles, The eukaryotic replisome tolerates leading-strand base damage by replicase switching, EMBO J 40 (2021) e107037. 10.15252/embj.2020107037.

[58] N. Sabouri, E. Johansson, Translesion Synthesis of Abasic Sites by Yeast DNA Polymerase ϵ, J. Biol. Chem. 284 (2009) 31555–31563. 10.1074/jbc.M109.043927.

[59] J.S. Lewis, L.M. Spenkelink, G.D. Schauer, O. Yurieva, S.H. Mueller, V. Natarajan, G. Kaur, C. Maher, C. Kay, M.E. O’Donnell, A.M. van Oijen, Tunability of DNA Polymerase Stability during Eukaryotic DNA Replication, Molecular Cell 77 (2020) 17-25.e5. 10.1016/j.molcel.2019.10.005.

[60] L.I. Toledo, M. Altmeyer, M.-B. Rask, C. Lukas, D.H. Larsen, L.K. Povlsen, S. Bekker-Jensen, N. Mailand, J. Bartek, J. Lukas, ATR Prohibits Replication Catastrophe by Preventing Global Exhaustion of RPA, Cell 155 (2013) 1088–1103. 10.1016/j.cell.2013.10.043.

[61] L. Toledo, K.J. Neelsen, J. Lukas, Replication Catastrophe: When a Checkpoint Fails because of Exhaustion, Molecular Cell 66 (2017) 735–749. 10.1016/j.molcel.2017.05.001.

[62] R.E. Georgescu, L. Langston, N.Y. Yao, O. Yurieva, D. Zhang, J. Finkelstein, T. Agarwal, M.E. O’Donnell, Mechanism of asymmetric polymerase assembly at the eukaryotic replication fork, Nat Struct Mol Biol 21 (2014) 664–670. 10.1038/nsmb.2851.

[63] M.R. Wasserman, G.D. Schauer, M.E. O’Donnell, S. Liu, Replication Fork Activation Is Enabled by a Single-Stranded DNA Gate in CMG Helicase, Cell 178 (2019) 600-611.e16. 10.1016/j.cell.2019.06.032.

[64] J. Yin, A.J. Lin, D.E. Golan, C.T. Walsh, Site-specific protein labeling by Sfp phosphopantetheinyl transferase, Nat Protoc 1 (2006) 280–285. 10.1038/nprot.2006.43.

[65] Z. Zhou, P. Cironi, A.J. Lin, Y. Xu, S. Hrvatin, D.E. Golan, P.A. Silver, C.T. Walsh, J. Yin, Genetically encoded short peptide tags for orthogonal protein labeling by Sfp and AcpS phosphopantetheinyl transferases, ACS Chem. Biol. 2 (2007) 337–346. 10.1021/cb700054k.

[66] A.E. Shaw, M.N. Mihelich, J.E. Whitted, H.J. Reitman, A.J. Timmerman, M. Tehseen, S.M. Hamdan, G.D. Schauer, Revised mechanism of hydroxyurea-induced cell cycle arrest and an improved alternative, Proc. Natl. Acad. Sci. U.S.A. 121 (2024) e2404470121. 10.1073/pnas.2404470121.

[67] C. Joo, T. Ha, Preparing sample chambers for single-molecule FRET, Cold Spring Harb Protoc 2012 (2012) 1104–1108. 10.1101/pdb.prot071530.

[68] C.E. Aitken, R.A. Marshall, J.D. Puglisi, An Oxygen Scavenging System for Improvement of Dye Stability in Single-Molecule Fluorescence Experiments, Biophysical Journal 94 (2008) 1826–1835. 10.1529/biophysj.107.117689.

[69] M.F. Juette, D.S. Terry, M.R. Wasserman, R.B. Altman, Z. Zhou, H. Zhao, S.C. Blanchard, Single-molecule imaging of non-equilibrium molecular ensembles on the millisecond timescale, Nat Methods 13 (2016) 341–344. 10.1038/nmeth.3769.

[70] S. Preus, S.L. Noer, L.L. Hildebrandt, D. Gudnason, V. Birkedal, iSMS: single-molecule FRET microscopy software, Nat Methods 12 (2015) 593–594. 10.1038/nmeth.3435.

[71] V.V. Kushnirov, Rapid and reliable protein extraction from yeast, Yeast 16 (2000) 857–860. 10.1002/1097-0061(20000630)16:9%253C857::AID-YEA561%253E3.0.CO;2-B.

