## Supplementary Information for "DNA Polymerase α has pyrimidine dimer translesion activity that is suppressed during normal replication"

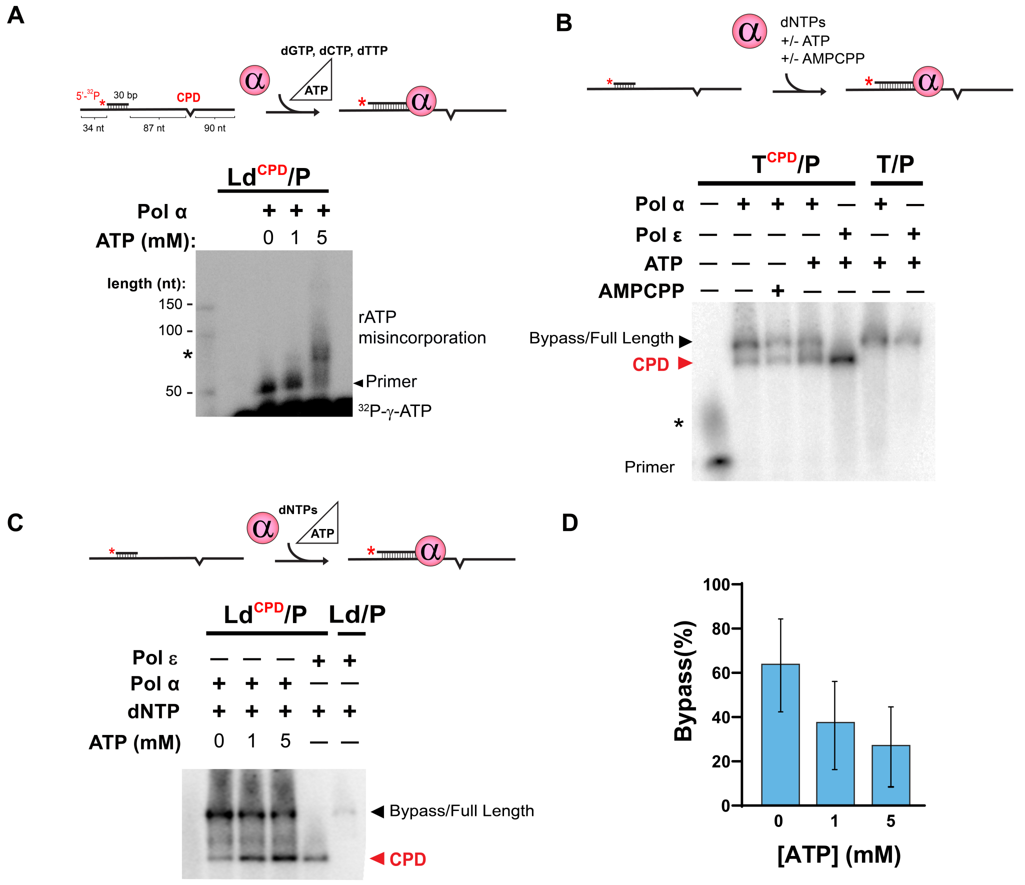


**Figure S1. ATP is moderately misincorporated into Pol α replication products when lacking dATP, but is inhibitory to TLS activity when all dNTPs are present. A)** A primed CPD template (Ld$^{CPD}$/P) is incubated with Pol α, a limited set of deoxynucleotides (dGTP, dCTP, dTTP), and increasing ATP concentrations. At 5 mM ATP, the majority of the product can be seen in the poly(dT) stall site upstream of the lesion (marked by asterisk). The bottom band is unincorporated $^{32}$P-γ-ATP, included to show both how high the contrast is turned up to visualize the small amount of product, and also how this particular gel ran askew, altering how the primers in the first two lanes run. **B)** (Ld$^{CPD}$/P) is incubated with Pol α, dNTPs, and the indicated nucleotide. AMPCPP is an ATP analog with a nonhydrolyzable γ phosphate. Pol ε is used as an internal control, stalling at the CPD, and the Ld/P substrate is used to show both Pols reaching full length. **C)** Representative denaturing gel showing titration of ATP into a TLS reaction containing dNTPs. Pol ε is included for Ld$^{CPD}$/P and Ld/P substrates to respectively show CPD-stalled and full length replication products. All lanes in A-C were stopped at 30 min. **D)** Mean and STD of the bypass percentage, or the ratio of full length product to CPD product intensity, quantified from three separate experiments like the one shown in panel C.

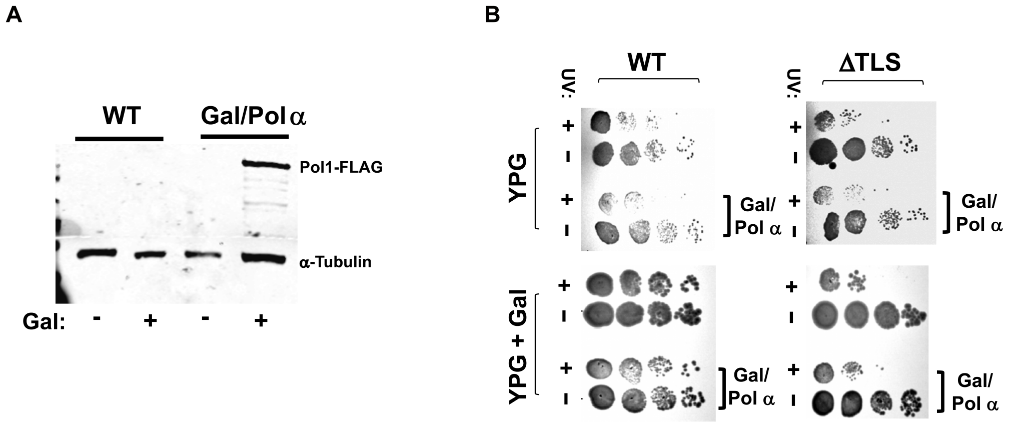


**Figure S2. Overexpression of Pol α does not rescue UV sensitivity in cells lacking TLS Polymerases. A)** Western blot of Pol1-FLAG in uninduced vs induced cell extracts. WT strain or Gal/Pol α overexpression strain (GAL/POL1-FLAG; GAL/PRI2; GAL/PRI2; GAL/POL12) were grown in the presence or absence of 2% Galactose as indicated, lysed, blotted by mouse anti-flag antibodies, and visualized by fluorescent anti-mouse secondary antibody, with a-Tubulin as a loading control. **B)** Yeast survival assays demonstrate that inducible Pol α overexpression does not rescue UV sensitivity in ΔTLS cells. Strains with a WT or ΔTLS background were integrated with all four Pol α genes under galactose control where indicated (Gal/Pol α). Strains were grown in Yeast Peptone Glycerol (YPG) and were incubated with 2% galactose where indicated. Where indicated, strains were exposed to 150 J/m$^2$ UV-C light, and strains under all conditions were then plated in a yeast survival spot assay (see Methods).

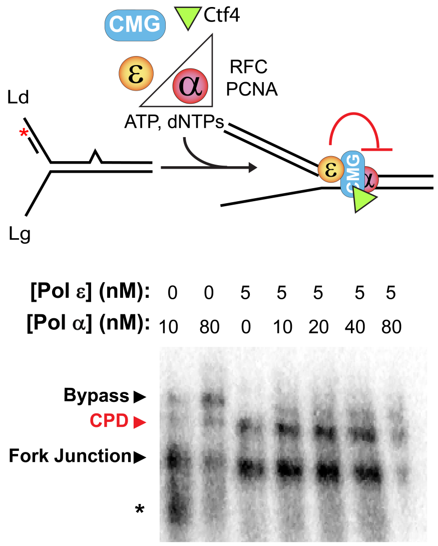


**Figure S3. Pol ε strongly inhibits Pol α TLS regardless of presence of Ctf4**
**A)** CMG is loaded onto forked CPD template. Pols α and/or ε are introduced simultaneously at the indicated concentration, in the presence of 120 nM Ctf4. The reaction is initiated with ATP and dNTPs. As in Fig 3, A small amount of Pol ε inhibits Pol α CPD TLS activity and is not outcompeted by excess Pol α. Note that much of Lane 7 diffused off the gel, but the lack of significant bypass is still apparent.

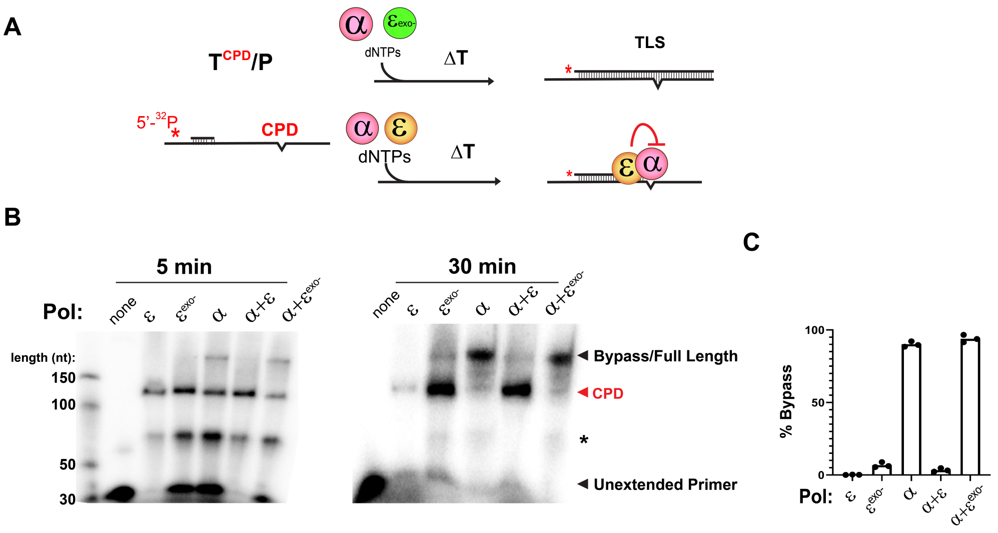


**Figure S4. Pol ε inhibits Pol a CPD TLS activity via exonuclease proofreading.**
**A)** Schematic of experiment. Pol e or Pol $\varepsilon^{exo-}$ (lacking 5'-3' proofreading ability) is added to Pol α CPD TLS reactions. **B)** Primer extension assays of two timepoints of Pol ε inhibition of TLS are shown, with the indicated Pols added simultaneously. During the first 5 minutes, there is little CPD bypass and much of the extended primer is stuck at the poly(dT) region in the template denoted by asterisk. After 30 min, both Pol ε and Pol $\varepsilon^{exo-}$ remain stuck at the lesion (with a small amount of full length product for Pol $\varepsilon^{exo-)}$, and Pol α exhibits efficient TLS activity. In both cases, Pol ε strongly inhibits Pol a TLS, whereas Pol $\varepsilon^{\varepsilon xo-}$ does not. **C)** The results were quantified for three separate 30 min timepoints. The bars indicate the mean of the individual values measured, which are shown as black circles.

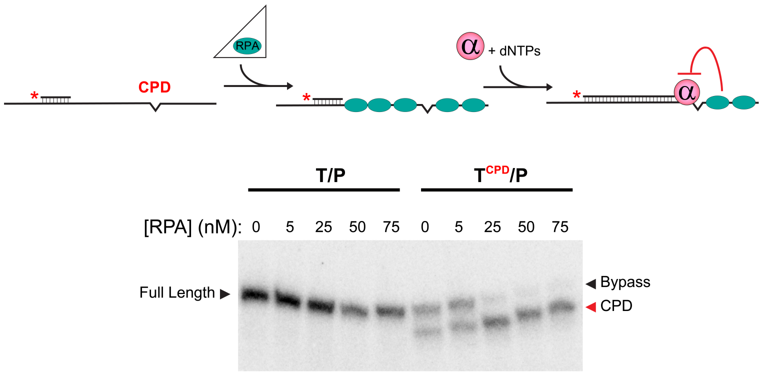


**Figure S5. RPA inhibits Pol α TLS activity on ssDNA substrates.**
The primed ssDNA substrate is coated with the indicated amount of RPA, followed by incubation with Pol α and dNTPs. Denaturing PAGE results of primer extension in the absence of presence of a CPD lesion, as indicated.

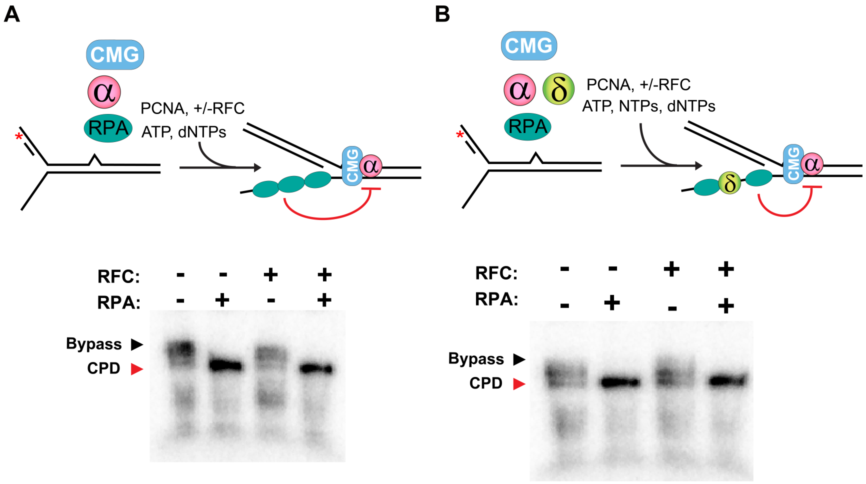


**Figure S6. RPA inhibits Pol α TLS activity at the replicaiton fork regardless of the presence or absence of RFC or Pol δ.** The reaction scheme in both **A)** and **B)** takes place as in Fig 2B, but 50 nM RPA is added in the replication initiation mix along with dNTPs and ATP where indicated, and 10 nM of RFC is added where indicated. In panel **B**, the presence of Pol δ and NTPs was included.

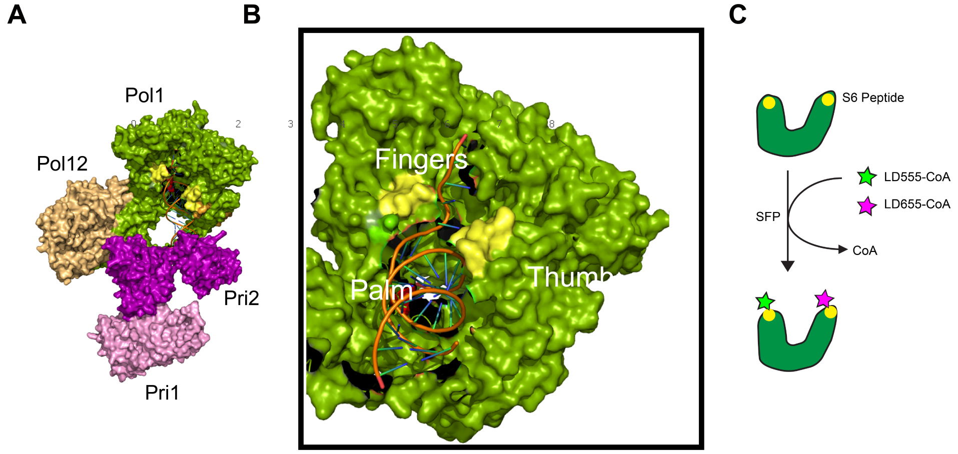


**Figure S7. Thumb and fingers labeling strategy to probe Pol α polymerase binding cleft conformation.**

**A)** The Pol α holoenzyme, with each subunit colored and labeled. The location of the solvent-accessible loops in the thumb and fingers domains that were inserted with the S6 peptide tag for labeling (see Methods) are colored in yellow. Figure rendered in Pymol with a cryoEM structure of Pol α in DNA elongation mode from Yuan et al., 2023. (PDB ID 8FOK). **B)** A closeup view of Pol1, showing the thumb and fingers engagement on the template. **C)** Schematic of SFP labeling. The S6 peptide was inserted into the thumb and fingers domains, and the SFP enzyme was used to covalently label these sites with CoA-derived fluorophores (LD555 or LD655). Using this approach, a fraction of the labeling reaction will result in a Pol α labeled with a FRET pair at the thumb and fingers domains.

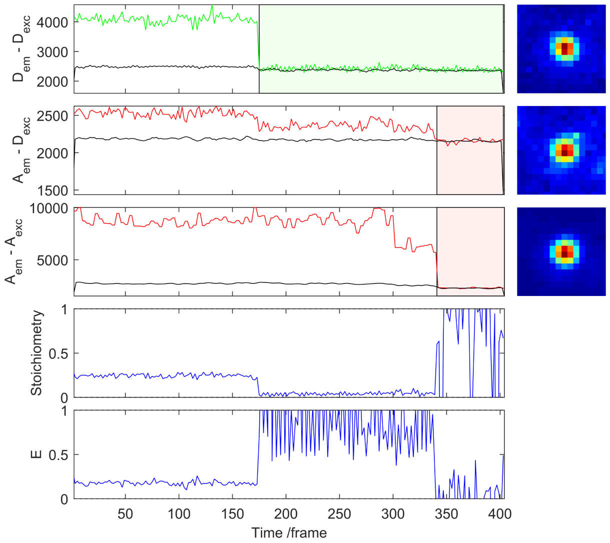


**Figure S8. Example of single-molecule fluorescence trace filtering.**
Representative fluorescence intensity trace, automatically filtered in the ALEX routine of iSMS to remove potentially spurious data. From top down, the trajectory and point spread function heat map (if applicable) is shown for 1) Donor excited ($D_{ex}$) Donor emission ($D_{em}$); ($D_{ex}/D_{em}$), 2) Dex/Aem (i.e., from FRET), 3) Aex/Aem (i.e., ALEX trace), 4) D:A stoichiometry as in Figure S3.3, and 5) FRET E. iSMS removes post-photobleaching data highlighted in green and red. The point spread functions in each of the three separately probed fluorescence schemes show a clear signal from a single point emitter (i.e., a single molecule) absent of background from neighboring molecules. In this trajectory, low FRET is verified by the existence of a donor bleaching concomitant with a drop in $D_{ex}/A_{em}$. The existence of the Aex/Aem (ALEX) signal that ultimately bleaches is further confirmation that a FRET pair was present.

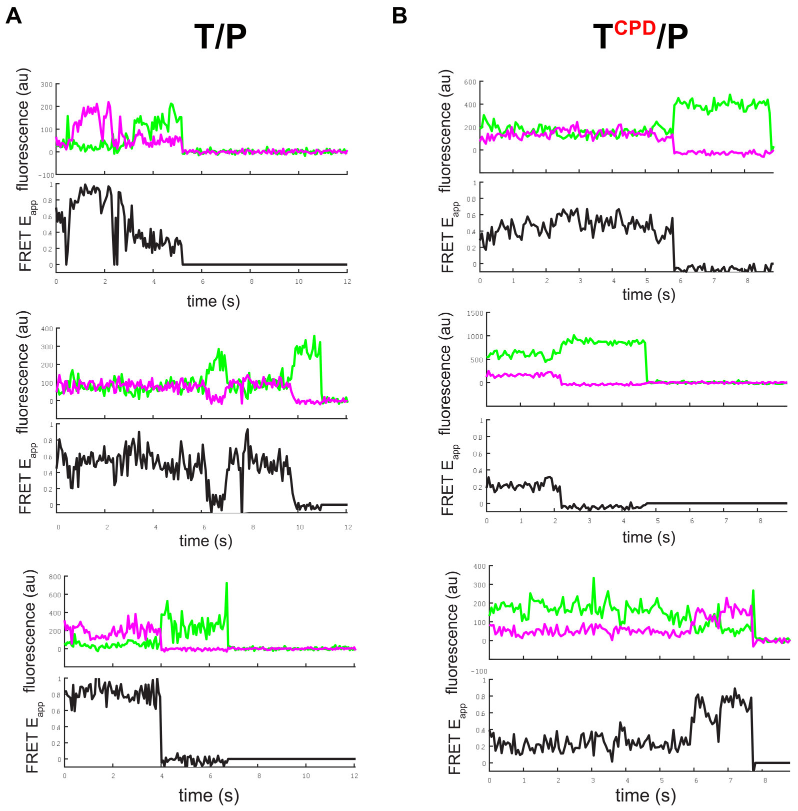


**Figure S9. Additional FRET traces from the histograms in Figure 3.**
Shown are acceptor-then-donor photobleach events from the histograms shown in Figure 3 for **A)** the regular template/primer or **B)** the CPD-template/primer. Note that low FRET traces like the ones observed at a CPD as (i.e., the main peak in Figure3D) will only rarely lead to acceptor photobleaching; these traces are not necessarily representative of the population but rather demonstrate the presence of dynamic and unambiguous FRET between the thumb and fingers domains during the experiment.

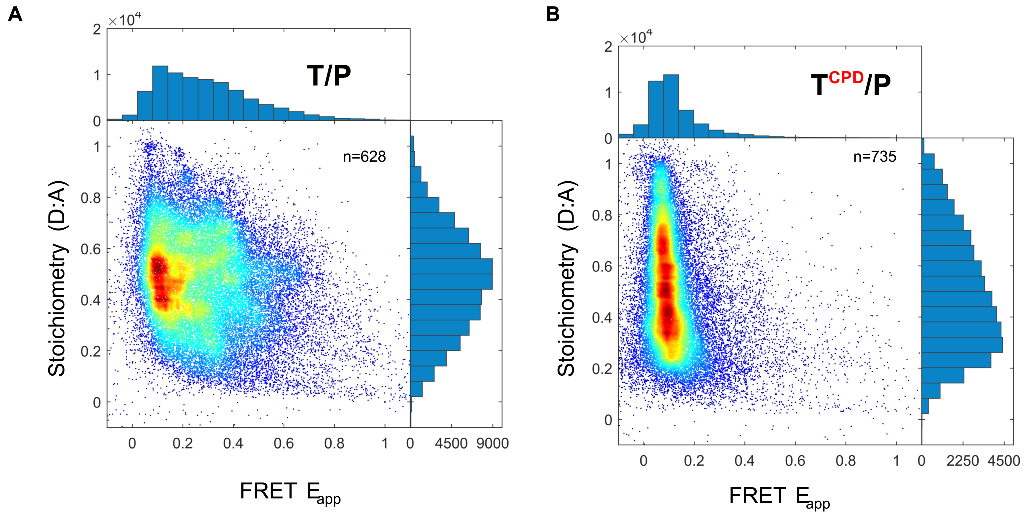


**Figure S10. Fluorophore stoichiometry of the histograms in Figure 3.**
Shown are the histograms from Figure 3 with an added dimension of dye stoichiometry, or ratio of D:A fluorophores for Pol a incorporating into the **A)** T/P or **B)** $T^{CPD}$/P substrate. Acceptable values for verifiable FRET typically fall within ~0.25 to 0.75. Plots Generated with iSMS.

**Table S1: Oligonucleotides used in this study**

| **Construct** | **Sequence** |
|---|---|
| **T**<br>(primed ssDNA assays) | CTCAGTCGCGATATATATATAGGTGTAGATTAATGTGGTTAGAATAGGGATGTGGTAGGAAG<br>TGAGAATTGGAGAGTGTGTTTTTTTTTTTTTTTTTTTTTTTTTTTTTTTTTTTTTTTTAAAGGT<br>GAGGGTTGGGAAGTGGAAGGATGGGCTCGAGACGTTTTTTTTTTTTTTTTTTTTTTTTTTTT<br>TTTTTTTTTTTT |
| **$T^{CPD}$**<br>(primed ssDNA assays) | CTCAGTCGCGATATATATATAGGTGTAGATTAATGTGG**<CPD>**AGAATAGGGATGTGGTAGG<br>AAGTGAGAATTGGAGAGTGTGTTTTTTTTTTTTTTTTTTTTTTTTTTTTTTTTTTTTTTTTTTAAA<br>GGTGAGGGTTGGGAAGTGGAAGGATGGGCTCGAGACGTTTTTTTTTTTTTTTTTTTTTTTT<br>TTTTTTTTTTTTTTTT |
| **P** | GTCTCGAGCCCATCCTTCCACTTCCCAACCCTCACC |
| **85DS_Ld**<br>(forked template assays) | AGAGAGTAGAGTGAGTTGTGGATGTGTAGAGTTGTTGTAGGAGAAGAGTTGTGAAGTGTG<br>GAGTGAGAGAAGAGAAGAGAGAGTGATATATTAATATTATATATTTTATAATTAATTAAATATAA<br>TTTTTTTTTTTTTTTTTTTTTTTTTTTTTTTTTTTTTTTTGAGGAAAGAATGTTGGTGAGGGTT<br>GGGAAGTGGAAGGATGGGCTTTTTTTTTTTTTTTTTTTTTTTTTTTTTTTTTTT |
| **85DS_Ld$^{CPD}$**<br>(forked template assays) | AGAGAGTAGAGTGAGTTGTGGATGTGTAGAGTTGTTGTAGGAGAAGAGTTGTGAAGTGTG<br>GAGTGAGAGAAGAGAAGAGAGAGTGATATA**<CPD>**AATATTATATATTTTATAATTAATTAAAT<br>ATAATTTTTTTTTTTTTTTTTTTTTTTTTTTTTTTTTTTTTTTTGAGGAAAGAATGTTGGTGAGG<br>GTTGGGAAGTGGAAGGATGGGCTTTTTTTTTTTTTTTTTTTTTTTTTTTTTTTTTTT |
| **85DS_Lg**<br>(forked template assays) | TTTTTTTTTTTTTTTTTTTTTTTTTTTTTTTTTTTTTTTTTTTTTTTTTTTTTTTTTTTTTTATATTT<br>AATTAATTATAAAATATATAATATTAATATATCACTCTCTCTTCTCTTCTCTCACTCCACACTTC<br>ACAACTCTTCTCCTACAACAACTCTACACATCCACAACTCACTCTACTCTCT |
| **85DS_P** | AGCCCATCCTTCCACTTCCCAACCCTCACC |
| **REV1 KO donor**<br>(template: gRNA plasmid #64329, marker: NTC). | <u>Forward:</u><br>TCAAAGAAGTAACGAGTTGACAGATTTTCTCAAAATAAATCGATACTGCATTTCTAGGCATA<br>TCCAGCGGACATGGAGGCCCAGAATACC<br><u>Reverse:</u>TCATGCAGATATTACAGGTAATGTTCGCAAACTGCGTGTTTACTGTATGCTGAAAT<br>GTTTTTTTTTTTTTAATTTAGGGGCAGGGCATGC |

| | |
|---|---|
| **REV3 KO donor** (template: pRS405, marker: LEU2): | Forward:<br>CAAAACAAGAGAAAGTATTTGAGTCAATACAAAACTACAAGTTGTGGCGAAATAAAAACTG<br>TGGGAATACTCAGGTATCGTAAGAT<br>Reverse:<br>AACGTTATACATAGAAACAAATAACTACTCATCATTTTGCGAGACATATCTGTGTCTAGATTA<br>AGCAAGGATTTTCTTAACTTCTTCG |
| **RAD30 KO donor** (template: pRS404, marker: TRP1): | Forward:<br>TAGTCTTCTAGCGCAGGCCTGCTCATTTTTGAACGGCTTTGATAAAACAAGACAAAGCATT<br>GTACTGAGAGTGCACCATAAACGACATTA<br>Reverse:<br>TATTATCAGGACGTTTTAGTTGCTGAAGCCATATAATTGTCTATTTGGAATACCTATTTCTTAG<br>CATTTTTGACGAAATTTGCTATTTTG |
| **Pol1$^{S6/S6}$ insertion donor** | TTATTAGCTAATCTGGTGGATCGTCGACGTGAAGTTAAGAAGGTGATGAAAACTGAAACTG<br>ATCCCCATAAGCGAGTTCAATGTGATATTCGTCAACAAGCTCTAAAATTGACTGCCAATTCT<br>ATGTATGGTTGTTTGGGTTATGTTAACAGTGGTGACTCCTTAAGTTGGTTGTTGAGATTGTT<br>GAACAGATTTTACGCAAAGCCGCTCGCCATGTTAGTCACTAATAAGGGTCGTGAGATTTTA<br>ATGAATACAAGACAACTAGCTGAAAGTATGAATCTTCTTGTAGTTTATGGTGATACAGATTC<br>GGTCATGATAGATACCGGTTGTGATAATTATGCGGATGCAATTAAAATTGGCTTGGGATTTA<br>AAAGGCTAGTAAATGAGCGCTATAGATTATTGGAGATTGATATTGATAATGTTTTTAAGAAGT<br>TACTATTACATGCCAAGAAAAAGTACGCTGCTTTGACTGTAAATTTGGACAAAAATGGTAAT<br>GGAACTACTGTTCTAGAAGTTAAAGGGTTGGACATGAAGCGTCGTGAATTTTGTCCACTTT<br>CGAGGGACGTTTCTATACATGTTTTAAACACCATCCTGTCAGATAAGGACCCAGAAGAAGC<br>ATTGCAAGAAGTGTATGATTACTTAGAAGACATCAGAATAAAAGTGGAAACCAATAACATTA<br>GAATTGATAAATATAAGATCAATATGAAGCTTTCAAAAGATCCCAAGGGAGATTCCTTAAGTT<br>GGTTATTAAGATTATTGAATGCCTACCCAGGTGGTAAAAACATGCCTGCAGTCCAAGTAGC<br>TCTAAGAA |
